# Genome-wide association study of school grades identifies a genetic overlap between language ability, psychopathology and creativity

**DOI:** 10.1101/2020.05.09.075226

**Authors:** Veera M. Rajagopal, Andrea Ganna, Jonathan R. I. Coleman, Andrea G. Allegrini, Georgios Voloudakis, Jakob Grove, Thomas D. Als, Henriette T. Horsdal, Liselotte Petersen, Vivek Appadurai, Andrew Schork, Alfonso Buil, Cynthia M. Bulik, Jonas Bybjerg-Grauholm, Marie Bækvad-Hansen, David M. Hougaard, Ole Mors, Merete Nordentoft, Thomas Werge, iPSYCH-Broad Consortium, Preben Bo Mortensen, Gerome Breen, Panos Roussos, Robert Plomin, Esben Agerbo, Anders D. Børglum, Ditte Demontis

## Abstract

Individuals with psychiatric disorders perform differently in school compared to the general population. Genetic factors contribute substantially to such differences. It is however unclear if differential performance is seen across all cognitive domains such as math and language. Here we report a genome-wide association study (GWAS) of school grades in 30,982 individuals (18,495 with and 12,487 without one or more of six major psychiatric disorders) and a replication study in 4,547 individuals. GWAS of overall school performance yielded results that were highly similar to the results of a previous GWAS of educational attainment. Analyzing subject specific grades, we observed that math performance was severely affected whereas language performance (Danish and English) was relatively unaffected or enhanced in those with psychiatric disorders compared to controls. We found that the genetic variants associated with poor math performance, but better language performance were also associated with increased risk for multiple psychiatric disorders. The same variants were also associated with creativity, which we show through a polygenic score analysis of 2953 creative professionals and 164,622 controls. The results overall suggest that risk for psychiatric disorders, language ability and creativity might have overlapping genetic roots.

## Introduction

Psychiatric disorders are common and have a complex etiology with contributions from both genetic and environmental factors.^1^ They are typically characterized by an early age of onset.^2^ While some disorders emerge during childhood, for example, autism spectrum disorder (ASD) and attention deficit hyperactivity disorder (ADHD), others emerge during adolescence or early adulthood, for example, schizophrenia (SCZ), bipolar disorder (BD) and anorexia nervosa (AN). Even long before the symptoms manifest, individuals often show signs of psychopathology.^3^ Several epidemiological studies have found atypical school performance as a risk factor for psychiatric disorders.^3^ Poor school performance has been shown as a risk factor for SCZ^4^ whereas excellent school performance for BD.^5^ Atypical school performance has been reported also in unaffected children and siblings of psychiatric patients suggesting that genetic factors could be involved.^3^

Large scale GWASs have been conducted for most of the major psychiatric disorders to date.^6–11^ Such studies have elucidated the complex genetic architecture of psychiatric disorders through identification of risk loci and wide-spread genetic correlations with a broad range of phenotypes. Through GWASs of psychiatric disorders it is now well established that a strong genetic overlap exists between many psychiatric disorders and cognitive phenotypes such as educational attainment and intelligence.^6–13^ For example, we have previously reported a strong negative genetic correlation between ADHD and educational attainment and a moderate positive genetic correlation between ASD and educational attainment in the largest GWASs of ADHD^7^ and ASD^6^ respectively. Also, a recent GWAS has reported a strong positive genetic correlation between AN and educational attainment.^9^ For most of the psychiatric disorders the genetic correlations with educational attainment align with the corresponding phenotypic associations with school performance reported in epidemiological studies.^14–16^ For some disorders such as SCZ, however, the genetic correlations do not correspond to the phenotypic associations.^17^ Although clinical and epidemiological studies have documented that individuals with—or at risk for—SCZ perform poorly in school^4^ and score low in neurocognitive assessments,^18^ the genetic correlation of SCZ with educational attainment is not negative, but rather positive, albeit weak.^17^ Similarly, although BD has been shown to be associated with cognitive deficits,^19^ it shows moderate positive genetic correlation with educational attainment.^8^ Such findings indicate that heterogeneity exists in the genetic overlap between the psychiatric disorders and cognitive phenotypes. The current large GWASs of cognitive phenotypes such as educational attainment^12^ and intelligence^13^ do not inform clearly about what causes such heterogeneities. GWASs of subject-specific school grades may help to study genetic associations of individual cognitive domains with psychiatric disorders, which in turn may help to disentangle the complex genetic overlap of psychiatric disorders with cognition. However, such analyses have not been possible so far, as the existing GWASs of school grades were based on small sample sizes and hence were not statistically powerful enough to study the genetic overlap with the psychiatric disorder.^20–23^

Here we present the largest GWAS of school grades to date disentangling the polygenic architecture of various domains of school performance and their complex phenotypic and genetic relationships with six major psychiatric disorders (ADHD, ASD, SCZ, BD, major depressive disorder (MDD) and AN). The overall study design is shown in Supplementary Fig. 1. Our discovery sample comes from iPSYCH^24^ and the Anorexia Nervosa Genetics Initiative (ANGI),^25^ large population-based Danish cohorts of individuals with and without psychiatric disorders for whom information on school grades was available through the Danish education register.^26^ Using principal component analysis (PCA), we decomposed the school grades in Danish, English and mathematics (six different grades per individual; Methods) into orthogonal principal components (hereafter, E-factors) that captured distinct cognitive domains relating to math and language. The GWASs of the E-factors (E1, E2, E3 and E4) identified multiple genome-wide significant loci. Among the E-factors, E1 correlated the most with educational attainment^12^ and intelligence.^13^ E2 captured differences between math and language performances and showed positive phenotypic and genetic correlations with almost all the psychiatric disorders thereby revealing a previously unknown relationship of psychiatric disorders with math and language cognitive domains. In addition, E2 showed a positive genetic association with creativity suggesting a shared genetic basis between language ability and creativity.

## Results

### Sample characteristics

Totally 30,982 individuals from iPSYCH^24^ and ANGI^25^ with information on school grades were included in our primary analysis after various quality control procedures (Methods). Among those, 18,495 (60%) had at least one of the six psychiatric disorders and 12,487 (40%) did not have any of the six disorders. The sample characteristics within each disorder and controls are shown in Table 1. It should be noted that the individuals included in our analysis were biased towards those with an overall better cognitive functioning as only such individuals manage to attend school and complete exams and hence have their grades recorded in the registers.^27^ The age of the study individuals (as of Dec 2016) ranged between 15 to 38 years (mean=24.2; SD=4.2). There were 14,606 males and 16,376 females. The school grades were from the exit exam (or ninth level exam or FP9) given at the end of compulsory schooling in Denmark. Age at the time of examination (hereafter, exam age) ranged between 14.5 and 17.5 years (mean=15.7; SD=0.42). Grades in three subjects namely Danish, English and mathematics were chosen since they were compulsory and so available for almost all individuals. Individuals with psychiatric diagnoses (as of Dec 2016) were considered as cases irrespective of whether the diagnoses were given before or after the exit exam.

**Table 1.** Sample characteristics

|  | **SCZ** | **BD** | **MDD** | **ADHD** | **ASD** | **AN** | **Controls** |
| --- | --- | --- | --- | --- | --- | --- | --- |
| **Sample size** | 1356 | 839 | 9719 | 5238 | 3859 | 1680 | 12487 |
| **Females** |  |  |  |  |  |  |  |
| *N* | 668 | 561 | 6836 | 1748 | 1017 | 1568 | 6327 |
| *%* | 49.3% | 66.9% | 70.3% | 33.4% | 26.4% | 93.3% | 50.7% |
| **Age** |  |  |  |  |  |  |  |
| *Mean* | 26.1 | 26.8 | 26.2 | 22.9 | 22.1 | 24.8 | 23.4 |
| *SD* | 2.9 | 2.8 | 3.3 | 4.1 | 3.9 | 3.7 | 4.2 |
| **Exam age** |  |  |  |  |  |  |  |
| *Mean* | 15.7 | 15.6 | 15.6 | 15.8 | 15.8 | 15.6 | 15.6 |
| *SD* | 0.4 | 0.4 | 0.4 | 0.4 | 0.5 | 0.4 | 0.4 |
| **Age at first diagnosis** |  |  |  |  |  |  |  |
| *Mean* | 21.15 | 21.86 | 18.9 | 14.8 | 12.8 | 16.3 | - |
| *SD* | 2.9 | 3.2 | 3.3 | 5.6 | 5.0 | 3.3 | - |
| **Diagnosed before exam** |  |  |  |  |  |  |  |
| *N* | 38 | 17 | 1655 | 3025 | 2880 | 819 | - |
| *%* | 2.8% | 2.1% | 17.1% | 57.8% | 74.6% | 48.9% | - |
| **School grades** |  |  |  |  |  |  |  |
| *Mean* | 6.0 | 6.8 | 6.4 | 5.3 | 6.7 | 8.0 | 6.9 |
| *SD* | 2.2 | 2.3 | 2.3 | 2.2 | 2.3 | 2.3 | 2.4 |

SCZ - Schizophrenia; BD - Bipolar disorder; MDD - Major depressive disorder; ADHD - Attention deficit/hyperactivity disorder; ASD - Autism spectrum disorder; AN - Anorexia nervosa
Age of the individuals on December 2016
Exam age is the age of the individuals when they sat for the exit exam.
The age at first diagnosis was calculated based on the first time the diagnosis was recorded in the register.
The date of first diagnosis and the date of exit exam were compared to identify if the individual has received the diagnosis before they sat for the exam.
The mean value was calculated across all the grades: Danish written, oral and grammar and English oral and mathematics written and oral (if exam year ≤ 2006) or problem solving with and without help (if exam year $>$ 2006); The lowest and highest mean values observed were -1.7 and 12.

### Decomposition of school grades in to cognitive domains

For each individual we had information on the following grades: Danish written, oral and grammar, English oral, mathematics written and oral (if sat for the exam before 2007), mathematics problem solving with help and problem solving without help (if sat for the exam after 2006; Methods). All the grades showed substantial heritability and were strongly correlated with each other both phenotypically and genetically (Supplementary Fig. 2). Individual GWASs of these grades may yield largely similar results and so may not be informative in studying the genetic overlap of distinct performance domains with the psychiatric disorders. Hence, we instead decomposed the grades into different cognitive domains using a principal component analysis (PCA). Since the mathematics exams were restructured in 2007, we performed PCA separately for grades given during 2002-2006 and 2007-2016 (Methods). We identified four informative E-factors that were reproducible between the two PCAs in terms of subject loadings and showed near perfect genetic correlations (rg ∼ 1) between the two datasets (Fig. 1a, 1b, 1c and Supplementary Table 1). After combining the two datasets, the four E-factors together explained 89.5% of the variance in the school grades (E1=56%; E2=13.5%; E3=10.5%; E4=8.5%). We discuss in detail how to interpret the four E-factors based on their subject specific loadings in the Supplementary Note. Briefly, E1 captured overall school performance (analogous to general cognitive ability factor^28^ (g) derived from a battery of cognitive tests), E2 captured relative differences between math and language performances, E3 captured relative differences between written and oral performances, and E4 captured relative differences between Danish and English performances (Table 2). We also repeated the PCA using only the control individuals (N=12,487), which yielded subject loadings largely similar to that of the main PCA (Supplementary Fig. 3). Hence, the subject loadings in the main PCA were not biased by including individuals with psychiatric disorders in the analysis.

**Figure 1.**
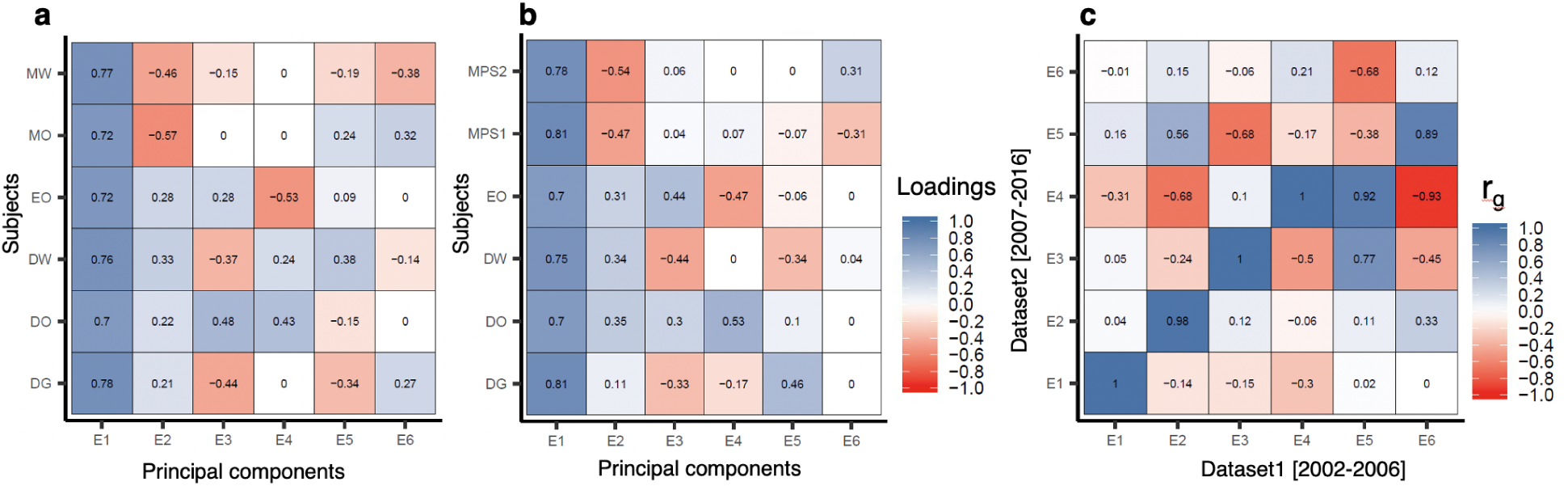
Loadings and genetic correlations of E-factors. **a**, PC loadings of the E-factors in dataset 1 [2002-2006; N=11,284]. **b**, PC loadings of the E-factors in dataset 2 [2007-2016; N=19,698]. **c**, Genetic correlations between E-factors from dataset 1 and dataset 2 calculated using GCTA bivariate GREML analysis; DG= Danish grammar; DO= Danish oral; DW= Danish written; EO=English oral; MO=Math oral; MW= Math written; MPS1= Math problem solving 1 (with help e.g. by using calculator); MPS2= Math problem solving 2 (without help)

**Table 2.** Description of E-factors

| E-factors | Variance | Description |
| --- | --- | --- |
| E1 | 56% | E1 captures overall school performance as the loadings were similar across all the subjects; the higher the E1 score, the better the individual in all the grades |
| E2 | 13.5% | E2 captures the difference between language and math performance; the higher the E2 score, the better the individual in language relative to math |
| E3 | 10.5% | E3 captures the difference between written and oral grades in Danish and English; the higher the E3 scores, the better the individual in oral exams relative to written exams |
| E4 | 8.5% | E4 captures the difference between English and Danish performances; the higher the E4 score, the better the individual in Danish relative to English |

The variance estimates represent the proportion of total variance in the school grades explained by the E-factors. The factors E2, E3 and E4 are relative measures, hence should be interpreted carefully. An individual can have a high E2 score even when the individual scored poorly in both language and math, if the individual's language grades are higher than their math grades

### GWAS of E-factors

We performed a GWAS for each of the four E-factors using a genetically homogenous sample of unrelated Europeans that comprised both individuals with and without psychiatric disorders (Methods). Psychiatric diagnoses as well as sex and exam age were included in the covariates as they were all significantly associated with the E-factors (Methods; Supplementary Table 2 and Supplementary Note).

We identified seven genome-wide significant loci, of which four were associated with E1, two with E2, one with E3 and none with E4 (Supplementary Table 3; Supplementary Fig. 4; Supplementary Dataset 1). Among these, only three remained strictly genome-wide significant (P<1.2×10^−8^) after adjusting for the four GWASs conducted. In a phenome-wide association study (see Methods) of the index variants in the seven loci, six were significantly associated with multiple cognitive phenotypes (Supplementary Table 4; Supplementary Dataset 2). Importantly, for three of the four loci identified for E1, educational attainment ranked highest among the 4,571 phenotypes tested. Hence, the genome-wide loci we identified are robust and are likely to be true positives. Biological annotations of the seven loci are discussed in the Supplementary Note. The common variants explained a significant proportion of the variance in all the four E-factors. The SNP-based heritability estimates were as follows: E1=0.29 (SE=0.01; P<1.0×10^−300^), E2=0.18 (SE=0.01; P<1.0×10^−300^), E3=0.13 (SE=0.01; P<1.0×10^−300^) and E4=0.08 (SE=0.01; P=1.0×10^−13^). We observed moderate levels of inflation in the GC lambda values, which were likely due to polygenicity rather than biases such as population stratification and cryptic relatedness as suggested by LD score regression analysis^29^ (Supplementary Table 5).

### Association of E-factors with educational attainment and intelligence

Among the four E-factors, E1 captured overall school performance hence we expected it to correlate well with educational attainment and intelligence in comparison to other E-factors. To evaluate this, we analyzed the genetic correlations of the E-factors with educational attainment^12^ and intelligence^13^ using LD score regression.^30^ As expected, among the four E-factors, E1 showed the highest genetic correlations with educational attainment (r_*g*_ =0.90; SE=0.03; P=4.8×10^−198^) and intelligence (r_*g*_ =0.80; SE=0.03; P=3.3×10^−128^) (Fig. 2a; Supplementary Table 6). The genetic correlation of E1 with educational attainment was significantly higher than that with intelligence (jackknife P=0.001; Methods). We also estimated polygenic scores for the iPSYCH cohort based on variant effect sizes from the GWASs of educational attainment and intelligence and tested their associations with the E-factors. The polygenic scores were strongly associated with E1 compared to other E-factors and explained 8.3% (educational attainment) and 4.9% (intelligence) of the variance in E1 (Fig. 2b, 2c; Supplementary Table 7). Overall the results suggested that E1, which measured the overall school performance, is slightly more comparable to educational attainment than to intelligence. The genetic correlations and polygenic score associations of E2, E3, E4 with years of education and intelligence were only modest compared to those of E1 (Fig. 2; Supplementary Tables 6, 7); the results are discussed in the Supplementary Note.

**Figure 2.**
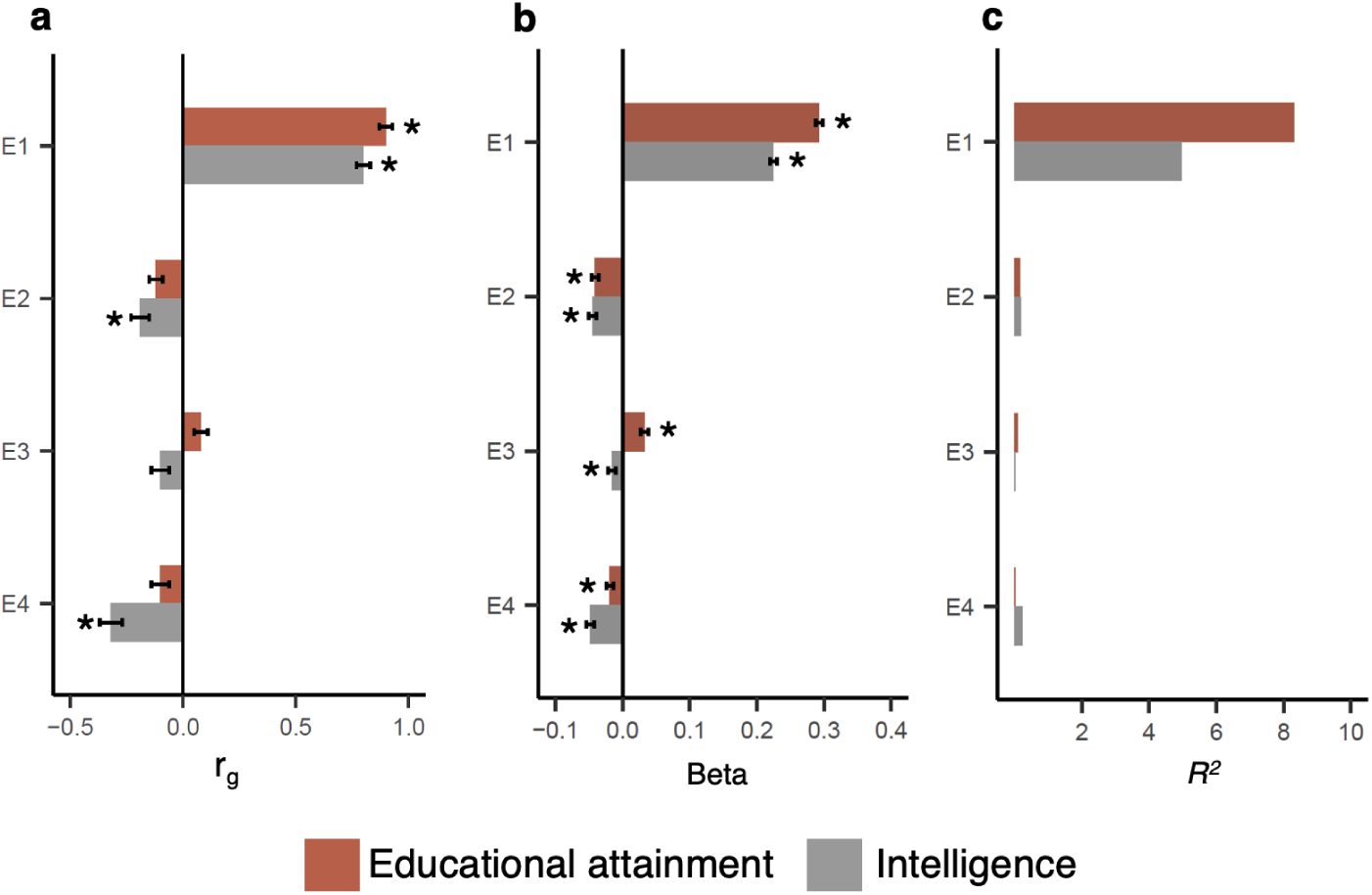
Association of E-factors with educational attainment and intelligence. **a**, Genetic correlations of E-factors with educational attainment and intelligence estimated using bivariate LD score regression **b**, Association of polygenic score for educational attainment and intelligence with E-factors (N=30,982; betas and standard errors are plotted). **c**, Maximum variance explained (R^2^) by the polygenic scores for years of education and intelligence in the E-factors. Star symbol indicates that the association is statistically significant after multiple testing correction (P<0.006)

### Association of E-factors with psychiatric disorders

In order to dissect the complex genetic overlap of school performance with psychiatric disorders, we evaluated the phenotypic and genetic associations of the E-factors with six psychiatric disorders (ADHD, ASD, SCZ, BD, MDD and AN). Phenotypic associations were evaluated by comparing the E-factor scores between each psychiatric disorder group and the controls. Genetic associations were evaluated using two approaches. First, we studied the genetic correlations of the E-factors with the six psychiatric disorders using LD score regression. Second, we constructed polygenic scores for the six psychiatric disorders in the iPSYCH cohort using variant effect sizes from recent large GWASs^6–11^ and then studied their associations with the E-factors only in the controls.

Among the E-factors, E1 (overall school performance) showed the strongest phenotypic associations with the psychiatric disorders (Fig. 3a; Supplementary Table 8). The factor E1 was positively associated with ASD, BD and AN, and negatively with SCZ, MDD and ADHD. The directions of the associations were in line with previous studies on the association of school performance with risk for psychiatric disorders.^4,5,14–16,31^ The genetic correlations of E1 with the psychiatric disorders mirrored the corresponding phenotypic associations in all except SCZ (Fig. 3b; Supplementary Table 9). The results suggested that the observed phenotypic associations are partly due to significant sharing of common risk variants between E1 and psychiatric disorders. In the case of SCZ, although E1 was significantly lower in cases compared to controls (Beta=-0.19; SE=0.02; P=1.4×10^−11^), the genetic correlation of E1 with SCZ was minimal (r_*g*_ =0.06; SE=0.03; P=0.06). Analysis of polygenic scores showed that all the six psychiatric disorders were associated with E1 in the controls in the same directions as the corresponding genetic correlations, which suggested that our genetic correlation results were less likely to be confounded by including psychiatric cases in the GWAS of E1 (Fig. 3c; Supplementary Table 10). However, the associations of E1 with the polygenic scores for SCZ (Beta=0.02; SE=0.008; adjusted P=0.09) and ASD (Beta=0.03; SE=0.01; adjusted P=0.07) were not statistically significant after multiple testing correction.

**Figure 3.**
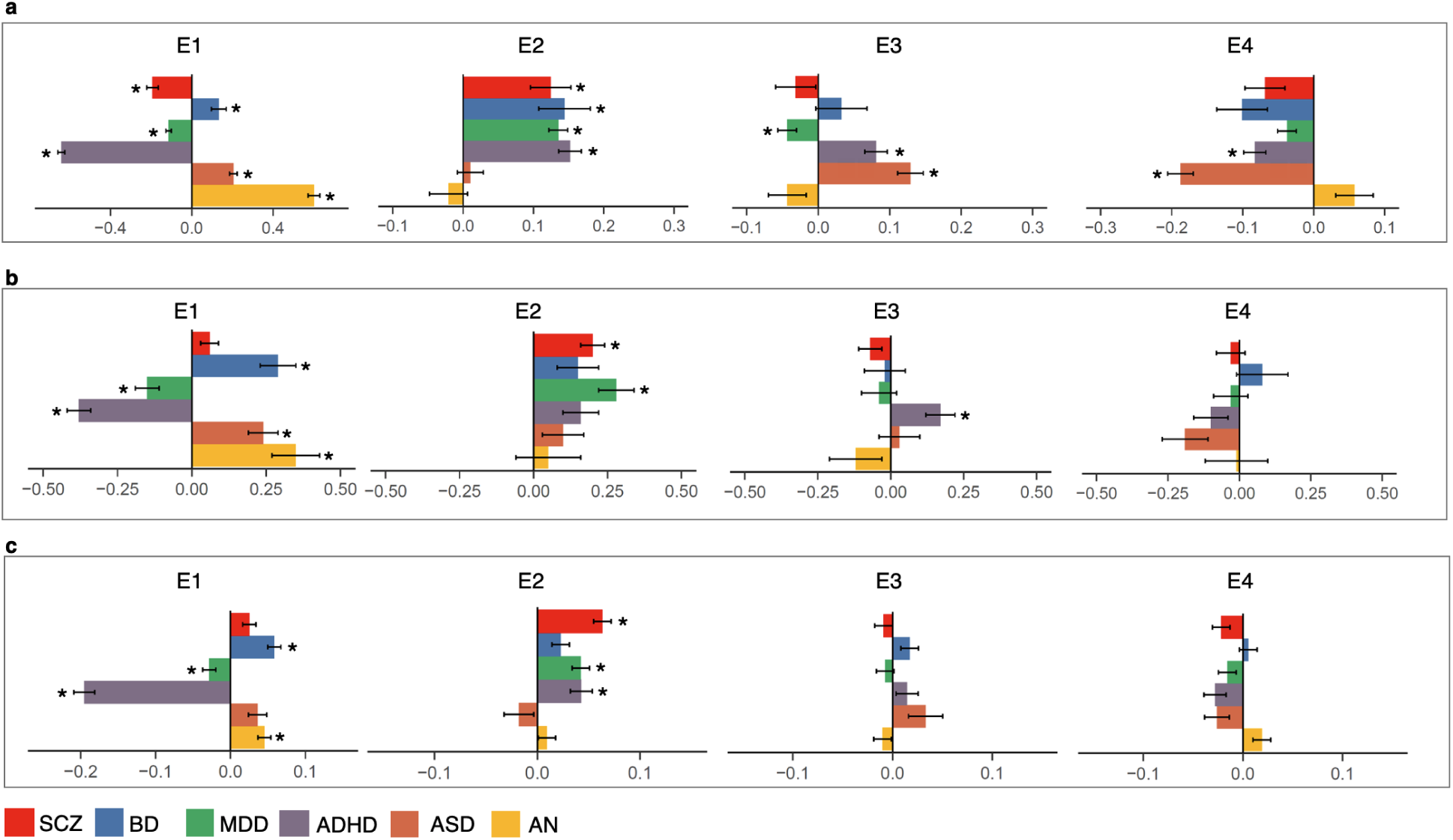
Association of E-factors with psychiatric disorders. **a**, Phenotypic associations of E-factors with six psychiatric disorders estimated using logistic regression (betas and standard errors are plotted). **b**, Genetic correlations of E-factors with six psychiatric disorders estimated using bivariate LD score regression (r_*g*_ and standard errors are plotted). **c**, Association of polygenic scores for six psychiatric disorders with E-factors only in control samples using linear regression (N=12,487; betas and standard errors are plotted). Star symbol indicates that the association is statistically significant after multiple testing correction (P<0.002); SCZ - Schizophrenia; BD - Bipolar Disorder; MDD - Major Depressive Disorder; ADHD - Attention Deficit Hyperactivity Disorder; ASD - Autism Spectrum Disorder; AN - Anorexia Nervosa

The factor E2 (language performance relative to math) was significantly associated with all the psychiatric disorders except ASD and AN (Fig. 3a; Supplementary Table 8). The E2 mean scores were significantly higher in ADHD, SCZ, BD and MDD compared to controls (meaning cases performed better in language relative to math compared to controls). Similar to the phenotypic associations, we observed positive genetic correlations for E2 with ADHD, SCZ, BD and MDD (Fig. 3b; Supplementary Table 9). The genetic correlations were however statistically significant only with SCZ (r_*g*_ =0.20; SE=0.04; P=2.1×10-5) and MDD (r_*g*_ =0.28; SE=0.06; P=1.1×10-5). We then tested the associations between polygenic scores for the six psychiatric disorders and E2 only in the controls and found similar results as the genetic correlations. The polygenic score associations were statistically significant for SCZ, MDD and ADHD (Fig. 3c; Supplementary Table 10). Overall, E2 showed positive associations with four of the six psychiatric disorders. The results suggested that individuals with SCZ, MDD, BD and ADHD performed significantly differently in language compared to math and the differential performance was at least partly due to sharing of common risk variants between these four disorders and E2.

The factors E3 (written performance relative to oral) and E4 (Danish performance relative to English) explained small amounts of variances in the school grades and did not show strong associations with the psychiatric disorders (Fig. 3; Supplementary Tables 8, 9, 10). The few statistically significant associations observed are discussed in the Supplementary Note.

### Association of language and math performances with psychiatric disorders

To further clarify the relationship of E2 with the psychiatric disorders, we compared the actual math and language grades between cases and controls. We calculated the mean across all math grades and the mean across all Danish and English grades and used them as measures of math and language performances respectively. Using a multiple logistic regression analysis (adjusted for exam age and sex), both math and language mean grades were included in the same regression model, thereby testing if the language performance differed between cases and controls after adjusting for the differences in the math performance and vice versa.

For all the six disorders, we observed substantial differences between math and language performances in cases compared to controls (Fig. 4a; Supplementary Table 11). At the phenotypic level, math grades were significantly lower in cases compared to controls for all the disorders except AN (Fig 4a; Supplementary Table 11). Of all the disorders, the SCZ cases had the lowest math grades compared to controls (Beta=-0.44; SE=0.03; P=3×10^−33^). Unlike math, the language grades were significantly higher in cases compared to controls in BD (Beta=0.24; SE=0.04; P=8.5×10^−8^), ASD (Beta=0.15; SE=0.02; P=2×10^−11^) and AN (Beta=0.35; SE=0.03; P=7.4×10^−25^). For ADHD, the language grades (Beta=-0.18; SE=0.02; P=1×10^−17^) were lower in cases compared to controls, though the difference was only less than half of the math difference (Beta=-0.52; SE=0.02; P=2.4×10^−123^). For SCZ (Beta=0.01; SE=0.03; P=0.64) and MDD (Beta=0.04; SE=0.01; P=0.006; adjusted P=0.07), no statistically significant differences were seen in the language grades between cases and controls.

**Figure 4.**
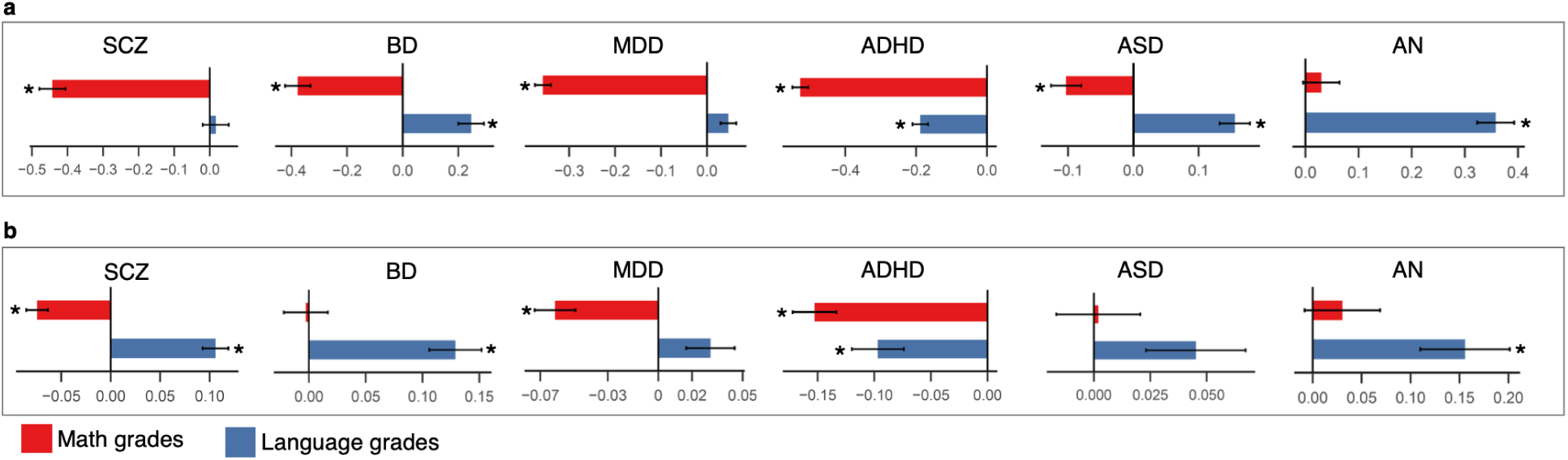
Association of math and language grades with psychiatric disorders. **a**, Phenotypic associations of math and language grades with six psychiatric disorders using multiple logistic regression (betas and standard errors are plotted). **b**, Association of polygenic scores for the six psychiatric disorders with math and language grades only in the control samples using multiple linear regression (N=12,487; betas and standard errors are plotted); Star symbol indicates that the association is statistically significant after multiple testing correction (P<0.004)

Next, we asked if language-math differences in the controls could also be influenced by genetic variants associated with psychiatric disorders. To test this, we studied the associations of polygenic scores for the six psychiatric disorders with language and math grades only in the controls using multiple linear regression analysis in the same way as we described earlier (covariates were same as those used in the main GWAS, except psychiatric diagnoses; see Methods). We found that the polygenic score associations were in the same directions as the corresponding phenotypic associations. The results suggested that even individuals who were not diagnosed with psychiatric disorders, but who had a higher polygenic risk for the psychiatric disorders, performed better in language relative to math (Fig. 4b; Supplementary Table 12). Hence, the language-math differences between psychiatric cases and controls were at least partly due to shared common variants influencing both math and language performances and risk for psychiatric disorders.

With regard to SCZ and BD, we observed an inverse association pattern between phenotypic analyses and polygenic score analyses (Fig. 4a, 4b). The SCZ cases had significantly poorer math grades, but not better language grades compared to controls (math: Beta=-0.44; SE=0.03; P=3×10^−33^; language: Beta=0.01; SE=0.03; P=0.64). However, individuals without SCZ, but with a higher polygenic risk for SCZ had significantly poorer math grades as well as better language grades (math: Beta=-0.07; SE=0.01; P=1.4×10^−11^; language: Beta=0.10; SE=0.01; P=5.1×10^−16^). This pattern was inverse for BD. The BD cases had significantly poorer math grades as well as better language grades compared to controls (math: Beta=-0.37; SE=0.04; P=1×10^−16^; language: Beta=0.24; SE=0.04; P=8.5×10^−8^). However, individuals without BD, but with a higher polygenic risk for BD had better language grades, but not poorer math grades (math: Beta=-0.002; SE=0.01; P=0.88; language: Beta=0.12; SE=0.02; P=2×10^−8^). A similar inverse pattern between SCZ and BD has been reported previously with regard to genetic correlations with educational attainment and intelligence. SCZ shows significant negative genetic correlation with intelligence, but a small positive genetic correlation with educational attainment^17^ (also E1) whereas BD shows significant positive genetic correlation with educational attainment^8^ (also E1) but no genetic correlation with intelligence. The results points to a unique inverse relationship of SCZ and BD with cognition, though the molecular mechanisms underlying this relationship are unclear.

### Replication in TEDS

To replicate our findings, we analyzed 4,547 genotyped individuals from the Twins Early Development Study (TEDS) cohort^32^ for whom General Certificate of Secondary Education (GCSE) school grades in English, math and science were available. The profile of the school grades in TEDS was similar to the one in iPSYCH: school grades were from the end of compulsory schooling; individuals were aged 15-16 years at the time of the examinations; the school grades were available in both math and language exams.^33^

PCA of English, math and science grades in TEDS yielded subject loadings that were similar to the subject loadings in the iPSYCH: E1 (first PC) had similar positive loadings from all three subjects; E2 (second PC) had positive loading from English and negative loadings from math and science; importantly, the English (0.81) and math (−0.50) loadings were higher than science loading (−0.27) suggesting that E2 captured mainly math and language performances (Fig. 5a). The factors E1 and E2 explained 83.4% and 10.4% of the variance in the school grades in TEDS respectively. Since the grades were not broken down to written and oral exams or a grade in foreign language that was taken by everybody was not available (unlike Denmark, where English exam is compulsory hence taken by everybody), we could not derive factors in TEDS equivalent to E3 and E4 in the iPSYCH.

We performed GWASs of E1 and E2 in TEDS and tested their genetic correlations with E1 and E2 in iPSYCH. The factors E1 and E2 in the two cohorts correlated almost completely (E1: r_*g*_ =0.99; SE=0.13; P=3.3×10^−13^; E2: r_*g*_ =1.07; SE=0.77; P=0.16; Fig. 5b; Supplementary Table 13). The genetic correlation of E2 between the cohorts, however, was not statistically significant due to the relatively smaller sample size of the TEDS. Nevertheless, when we predicted E1 and E2 in TEDS using polygenic scores (constructed using the effect sizes from GWASs of E1 and E2 in iPSYCH), we observed significant associations supporting the genetic correlation analyses (Fig. 5c; Supplementary Table 14).

**Figure 5.**
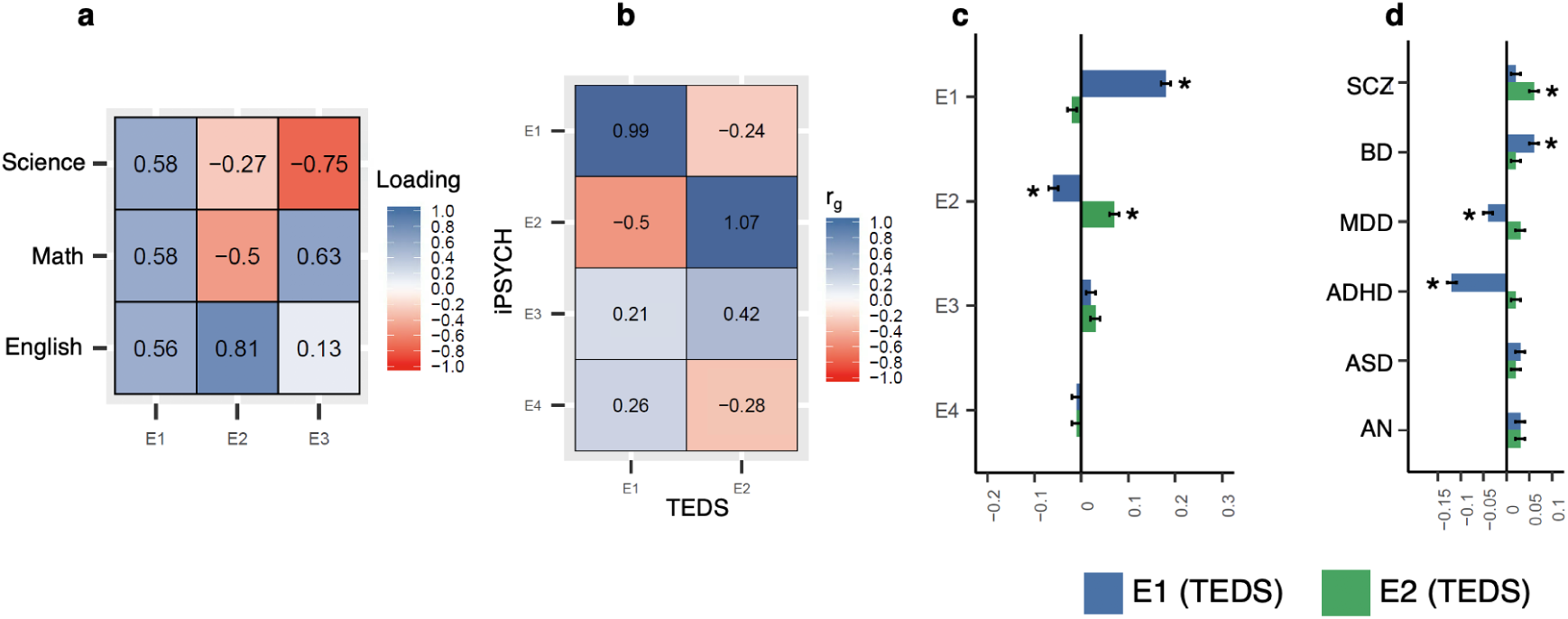
Replication analysis in TEDS. **a**, Subject loadings from PCA of school grades in TEDS. **b**, Genetic correlations of E1, E2, E3, and E4 in iPSYCH with E1 and E2 in TEDS estimated using bivariate LD score regression **c**, Association of polygenic scores for E1, E2, E3 and E4 in iPSYCH with E1 and E2 in TEDS using linear regression (N=4,547; betas and standard errors are plotted). **d**, Association of polygenic scores for psychiatric disorders with E1 and E2 in TEDS using linear regression (N=4,547; betas and standard errors are plotted). Star symbol indicates that the association is statistically signficant after multiple testing correction (P<0.006 for fig. c and P<0.004 for fig. d)

We also tested the genetic associations of E1 and E2 with the psychiatric disorders in TEDS using polygenic score analysis (Fig. 5d; Supplementary Table 15). The polygenic scores for the six psychiatric disorders were associated with E1 and E2 in TEDS in the same directions as seen in iPSYCH. Notably, we observed positive associations between the E2 and all the six psychiatric disorders, albeit most of the associations except SCZ were only borderline significant; SCZ showed the strongest association with E2 (Beta=0.06; SE=0.01; P=3.6×10^−6^). Hence, overall we observed an agreement in the results between iPSYCH and TEDS that the genetic variants associated with E2 (better performance in language relative to math) were also associated with increased risk for psychiatric disorders especially SCZ.

### Association of E2 with creativity

Creativity has been historically believed to relate positively with psychopathology.^34^ Many epidemiological studies^35–37^ and genetic studies^38,39^ have reported supporting findings. Our analyses showed that E2 was associated with increased risk for multiple psychiatric disorders as well as with increased language performance. Hence, we asked if common variants associated with E2 were also associated with creativity. To evaluate this we analyzed 167,575 individuals from the Million Veterans Program (MVP) biobank,^40^ for whom information on occupation category was available. We classified individuals employed in the category “arts, design, entertainment, sports and media” as “creative professionals” (N=2,953) and the rest as controls (N=164,622). We constructed polygenic scores for all four E-factors using effect sizes from the iPSYCH GWASs and compared the scores between the creative professionals and controls after adjusting for their highest level of education, age and sex along with other covariates (Methods). For sensitivity analysis, we also compared the scores of individuals in each of the other occupation categories against the rest. We found that the E2 polygenic scores were significantly higher in creative professionals compared to others. This association was specifically observed between E2 and creative occupation. That is, among the four E-factors, E2 showed the strongest association with creativity, and among 24 occupation categories, creative occupations showed the strongest association with E2. (Fig. 6; Supplementary Table 16). The results suggested that individuals with higher genetic predisposition to perform better in language relative to math in school (indexed by higher polygenic score for E2) have higher odds for being employed in a creative occupation later in life.

**Figure 6.**
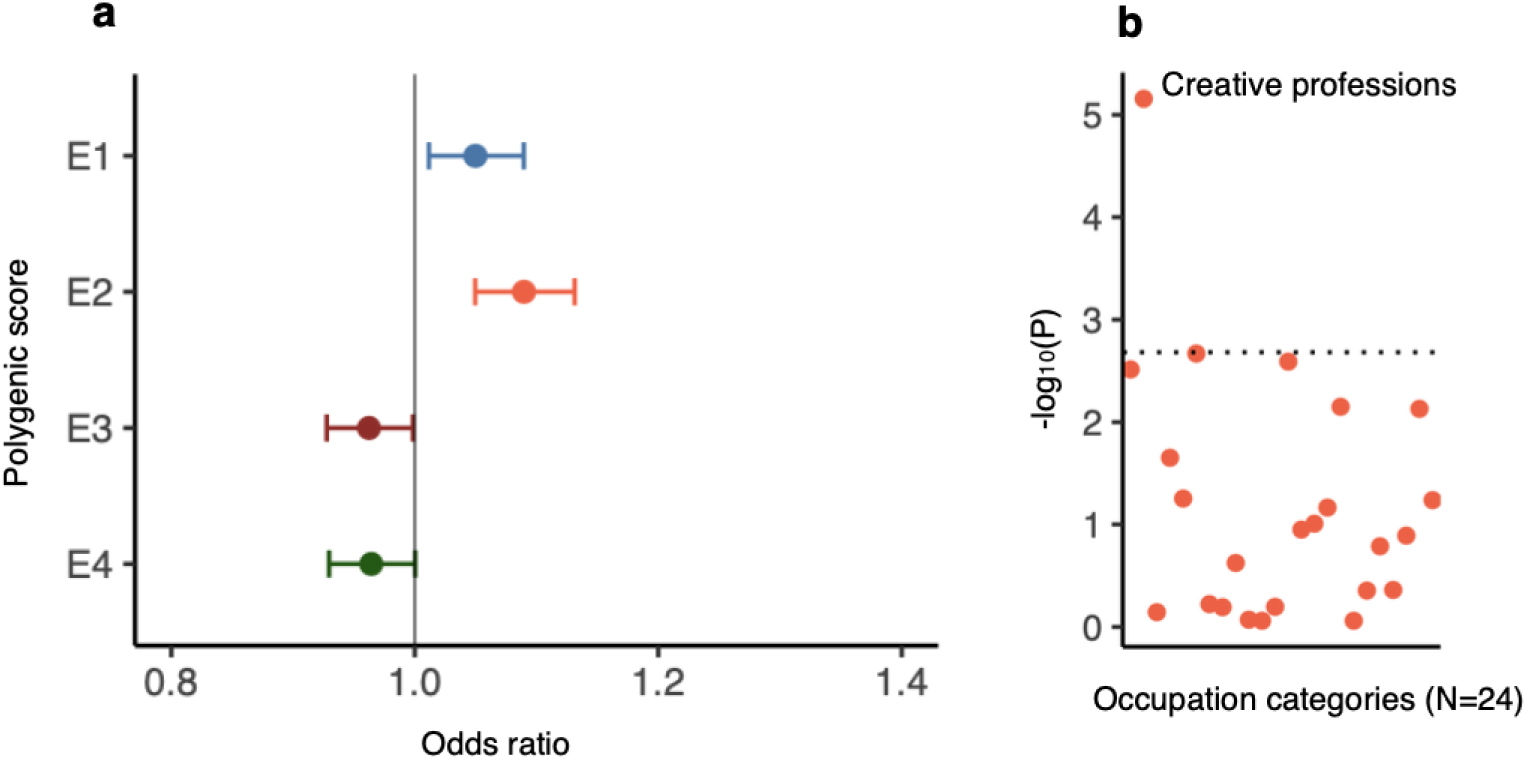
Association of polygenic scores for E-factors with occupation categories. **a**, Polygenic scores for E-factors were tested for association with creative profession (arts, design, entertainment, sports and media vs rest) using logistic regression; odds ratio and 95% confidence intervals are plotted. **b**, Polygenic score for E2 was tested for association with 24 occupation categories using logistic regression. Negative log_10_ P values are plotted. Dotted line correspond to P value threshold for significance after multiple testing correction (P<0.002)

## Discussion

To our knowledge, the results presented here represent the most comprehensive report to date of the phenotypic and genetic differences in subject specific school performances in individuals with and without psychiatric disorders. The study has been possible due to the unique register based resources linked to the iPSYCH^24^ and ANGI^25^ cohorts, which enabled us to combine genetic information of the study individuals with their school grades in the Danish education register.^26^ Our cohort is the largest available to date for the study of genetics of school grades. Our study poses notable advantages compared to previous GWASs of educational attainment.^12,41,42^ The phenotype, school grades, is fine grained hence capture more variance, for example, multiple individuals with the same number of years of education could differ substantially in their school grades. Our phenotype is objectively measured (graded by the teachers) unlike educational attainment, which is mostly self-reported hence likely to have more imprecision and heterogeneity compared to school grades. Our study participants were all from a single large cohort hence the study variables—school grades and psychiatric diagnoses—are likely to have less heterogeneity compared to meta-analytic studies where different cohorts follow different methods of ascertainment. Our study individuals were identified through nation-wide registers hence not subject to voluntary participation bias that has been shown to affect GWAS results particularly the genetic correlations.^43^

Availability of grades in multiple subjects for the same individuals enabled us to perform a PCA and decompose the school grades into distinct factors each representing unique cognitive abilities such as math and language. Due to a change in the Danish education system, the math exams were conducted differently since 2007. Hence, we had to split the dataset into two and perform PCA separately for data collected before and since 2007. However, it gave us an opportunity to internally replicate the PCA structure. The subject-specific loadings of the first four E-factors were remarkably similar between the two datasets. Moreover, we found near perfect (rg ∼ 1) genetic correlations between the corresponding E-factors in the two datasets suggesting that the common variant architectures of the corresponding E-factors were similar in the two datasets. We further replicated the PCA structure of E1 and E2 in an independent cohort, TEDS^32^ and demonstrated a near perfect genetic correlations for E1 and E2 between iPSYCH and TEDS. Hence, we have provided a compelling evidence that the signals captured by the E-factors, E1 and E2, are robust and reproducible.

The factor E1 measured the overall school performance and showed a strong genetic correlation with educational attainment,^12^ which suggested that most of the common variants that influence educational attainment also influence overall school performance. Importantly, both E1 and educational attainment showed similar genetic correlations with the six psychiatric disorders, which reinforced the conclusion that the common variant architecture is highly similar between overall school performance and educational attainment. Notably we observed that the genetic correlation between SCZ and E1 is positive, but minimal. Using polygenic score analysis, we further validated this finding by showing that the association between genetic liability to SCZ and E1 in the controls is also positive, but minimal. Our finding confirms similar earlier observations with regard to educational attainment.^17,44^ Hence, it is unlikely that the absence of a negative association between SCZ and educational attainment (as one would expect given that the individuals with SCZ perform poorly in school) in the previous studies were due to selection bias in the GWAS cohorts.^45^

The main findings of our study are those related to E2 as they offer novel insights into the relationship between the psychiatric disorders and cognition. The factor E2 measured the language performance relative to math performance and consistently showed positive phenotypic and genetic correlations with four of the six psychiatric disorders (SCZ, MDD, BD and ADHD). Further analysis of language and math grades confirmed that the positive correlations of E2 with the four psychiatric disorders are indeed due to increased language and decreased math performances in the cases compared to controls. Furthermore, language-math differences were also seen in ASD and AN, both of which did not show significant association with E2. Considering both the sets of analyses together, it is evident that heterogeneity in math and language cognition occurs in all the six psychiatric disorders, but the extent of the heterogeneity differs across the disorders. Our replication analysis supports our above interpretation. In the TEDS cohort, the polygenic scores for all the six psychiatric disorders showed positive associations with E2. However, only for SCZ the association was statistically significant. This is likely due to small sample size in TEDS. The fact that only SCZ showed significant association with E2 in both iPSYCH and TEDS suggests the heterogeneity in the math and language cognition is strongest for SCZ.

The E2 findings add two important insights. Firstly, the findings suggest that the positive genetic correlations of ASD,^6^ BD^8^ and AN^9^ with educational attainment^12^ were likely to be driven by language specific cognition as the language performance was better, but the math performance was poorer in cases compared to controls. Though individuals with higher genetic risk for ASD or BD or AN are genetically predisposed to attain higher education as suggested by the positive genetic correlation with educational attainment, we speculate that they do so by choosing a field that requires language skills rather than math skills. GWASs of educational attainment separately in language and math related fields in the future might help to confirm our assumption.

Secondly, the positive genetic association between E2 and risk for psychiatric disorders suggest that the genetic variants associated with increased risk for psychiatric disorders are either detrimental to math ability or beneficial to language ability or both. It is unclear if the positive link between psychiatric risk and language ability is a true biological link. If so, this association might have an evolutionary basis as has been suggested previously particularly in the context of schizophrenia.^46,47^ Alternatively, the association of psychiatric risk variants with increased language ability could be simply due to that the individuals with the risk variants compensate for their math deficits by being good in language.

Our final analysis suggested that individuals genetically predisposed to be relatively better in language than in math at school more often choose a creative occupation such as writing, acting and music later in life. As we have demonstrated that these individuals were also at risk for psychiatric disorders, the finding aligns with previous two studies that demonstrated that individuals with increased genetic risk for schizophrenia and bipolar disorder are more often involved in creative occupations^38^ or score better in creativity tests compared to general population.^39^ However, it is unclear if the association with creative occupations indicates that the individuals with the risk variants become creative due to their better language skills or simply choose creative professions as such professions suit well to their relatively poor math skills.

Our study has few limitations. First, our interpretations are based on the assumption that school performances in language and mathematics capture the cognitive abilities related to language and math domains respectively. However, our assumptions may not be entirely true as other factors including family^48^ and school socioeconomic statuses^49^ influence students’ school performances. Furthermore, like any other GWASs,^12,13^ the effect sizes that we measured in our study are likely to be inflated as we have not accounted for the rearing environment.^50^ Future studies that use a within-family design might help address these limitations. Second, although our primary sample, iPSYCH, has a better sampling design and do not suffer from participation bias,^43^ the individuals included in the current study are only a subset and are not representative of the full iPSYCH cohort.^24^ We included only those who were functional enough to go to school and attend the exams. Hence, our study sample is slightly biased towards better functioning individuals and as a result many of our findings cannot be generalized to the disorders, for example, findings related to ASD might apply only to the high functioning subtypes. Third, not all the individuals in the case groups received their diagnoses before the exams. This factor was not accounted for in our analysis since the diagnoses were register based and it is therefore not possible to confirm if the individuals were asymptomatic before the date when the diagnoses were first registered.

In summary, through an extensive analysis of subject specific school grades in a large sample, we have convincingly demonstrated for the first time that individuals with psychiatric disorders exhibit wide differences in their math and language cognitions. These differences seem to have a genetic basis understanding which is important as the knowledge may help to treat the cognitive deficits associated with these disorders.

## Acknowledgements

1. The iPSYCH project is funded by the Lundbeck Foundation (grant numbers R102A9118 and R155-2014-1724) and the universities and university hospitals of Aarhus and Copenhagen. The Danish National Biobank resource was supported by the Novo Nordisk Foundation. Data handling and analysis on the GenomeDK HPC facility was supported by NIMH (1U01MH109514-01 to Michael ODonovan and ADB). High-performance computer capacity for handling and statistical analysis of iPSYCH data on the GenomeDK HPC facility was provided by the Center for Genomics and Personalized Medicine, Aarhus University and Central Region Denmark, and Centre for Integrative Sequencing, iSEQ, Aarhus University (grant to ADB).
2. The Anorexia Nervosa Genetics Initiative (ANGI) was an initiative of the Klarman Family Foundation.
3. The PhD fellowship of V.M.R was fully funded by the Graduate School of Health, Aarhus University, Aarhus, Denmark.
4. G.V is supported by the Leon Levy Foundation (Leon Levy Fellowship in Neuroscience) and by NIH grant R01MH109677.
5. P.R is supported by the National Institutes of Health (R01AG050986 Roussos, R01MH109677 Roussos, U01MH116442 Roussos, R01MH110921 Roussos) and the Veterans Affairs (Merit grant BX004189 and BX002395 Roussos).
6. We gratefully acknowledge the ongoing contribution of the participants in the Twins Early Development Study (TEDS) and their families. TEDS is supported by a programme grant to RP from the UK Medical Research Council (MR/M021475/1 and previously G0901245), with additional support from the US National Institutes of Health (AG046938). The research leading to these results has also received funding from the European Research Council under the European Unions Seventh Framework Programme (FP7/2007-2013)/grant agreement n 602768 and ERC grant agreement n 295366.
7. R.P is supported by a Medical Research Council Professorship award (G19/2). This project has received funding from the European Unions Horizon 2020 research and innovation programme under the Marie Sklodowska-Curie grant agreement no. 721567.
8. A.G.A. has received funding from the European Union’s Horizon 2020 research and innovation programme under the Marie SklodowskaCurie grant agreement no. 721567
9. The authors acknowledge use of the research computing facility at Kings College London, Rosalind (https://rosalind.kcl.ac.uk), which is delivered in partnership with the National Institute for Health Research (NIHR) Biomedical Research Centres at South London & Maudsley and Guys & St. Thomas NHS Foundation Trusts, and part-funded by capital equipment grants from the Maudsley Charity (award 980) and Guys & St. Thomas Charity (TR130505). The views expressed are those of the author(s) and not necessarily those of the NHS, the NIHR, Kings College London, or the Department of Health and Social Care.

## Competing interests statement

1. CM Bulik reports: Shire (grant recipient, Scientific Advisory Board member); Idorsia (consultant); Pearson (author, royalty recipient).

## Author contributions

The author contributions are reported in Supplementary Table 19.

## Data availability

The GWAS summary statistics will be made available at https://ipsych.dk/en/research/downloads/.

## Online methods

### Brief summary of the cohorts involved in the study

Our study individuals come from four cohorts: iPSYCH,^24^ ANGI,^25^ TEDS^32^ and MVP.^40^ The main analyses were performed in iPSYCH and ANGI cohorts. iPSYCH is a large Danish case-cohort established for the study of genetic and environmental risk factors associated with major psychiatric disorders (ADHD, ASD, MDD, SCZ, BD).^24^ ANGI is an international collaboration established between scientists in United States, Australia/New Zealand, Sweden and Denmark to establish a cohort of individual with and without AN.^25^ The ANGI participants involved in our study were those recruited in Denmark along with the iPSYCH participants. Hence, they were QCed and genotyped together along with iPSYCH samples. All further descriptions about sample recruitment, genotyping and ethical approvals of iPSYCH cohort apply to ANGI as well. The phase-1 release^24^ of the iPSYCH cohort, after QC, comprises 77,639 individuals—51,101 with one or more of the six disorders and 27,605 controls—who were identified from a baseline sample comprising of the entire Danish population (N=1,472,762) born between 1981 and 2005. The cases were selected based on the psychiatric diagnosis information in the Danish Psychiatric Central Research Register^51^ and Danish National Hospital Register.^52^ The controls were randomly sampled from the baseline sample. In the current study, 30,982 genotyped individuals who had school grades information available in the Danish education register^26^ were included.

TEDS^32^ is a large twin cohort comprising of more than 16,000 twin pairs who were born either in England or Wales between 1994 and 1996. The twins were around 18 months old at the time of recruitment and were followed up longitudinally. Information on cognitive abilities, educational attainment, behavior and emotion were collected. Among the TEDS participants, 4547 individuals, comprising of one of each of the twin pairs, who were genotyped and also had information on their GCSE school grades were included in the current study.

The MVP^40^ is a large biobank at the Department of Veterans Affairs (VA), USA, established to study genetic and environmental influences on human diseases and traits. The participants being recruited into the study are active users of the US Veterans healthcare system. The MVP v3.0 data release comprised of 455,789 genotyped individuals. Information on demographics, health, lifestyle, medical history and family history were collected through questionnaires or through electronic health records in the US Veterans healthcare system.^40^ Among those in the MVP v3.0 release, 165,575 individuals, who were identified to be of European ancestries and provided information on occupation and educational attainment, were included in the current study.

### Ethical approvals

All the analyses in the iPSYCH data are within the permissions received from the Danish Scientific Ethics Committee, the Danish Health Data Authority, the Danish data protection agency and the Danish Neonatal Screening Biobank Steering Committee.^24^ The analyses in the TEDS cohort are within the permissions received from the Kings College London Ethics Committee (reference: PNM/09/10-104).^32^ Parental consent was obtained for all the TEDS participants before data collection. All the participants in the MVP cohort have given written informed consent at the time of recruitment.^40^ The analysis involving MVP data in the current study is approved by the VA Central Institutional Review Board.

### Ascertainment of psychiatric disorders in the iPSYCH cohort

The psychiatric diagnoses of the iPSYCH case samples were identified through Danish Psychiatric Central Research Register^51^ and Danish National Hospital Register.^52^ The diagnoses codes are based on International Classification of Diseases, 10th revision.^53^ The ICD-10 codes of the six disorders as follows: ADHD F90.0; ASD - F84.0, F84.1, F84.5, F84.8 or F84.9; MDD F32-39 [Since 96% of the individuals had either F32 (depressive episode) or F33 (recurrent depressive disorder), we call it as MDD rather than as affective disorders]; BD - F30-31; SCZ - F20; AN - F50.0. Individuals with mental retardation (ICD-10 F70-79) were excluded.

### School grades in the iPSYCH cohort

The school grades in the iPSYCH cohort were from the exit exam (also called as ninth level exam or FP9) given at the end of compulsory schooling in Denmark. The exam grades were obtained from the Danish education register^26^ that maintain school grades from all public schools in Denmark since 2001.

We chose three subjects namely Danish, English and mathematics for the current study. These subjects were chosen because they were compulsory, and so were available for a maximum number of individuals in the cohort. The examination types included written, oral and grammar in Danish, oral in English and either written and oral, or problem solving with and without help in mathematics. The data were from the exams conducted between 2002 and 2016.

The grades were in seven-point scale: −3 (unacceptable performance), 00 (inadequate performance), 02 (adequate performance), 4 (fair performance), 7 (good performance), 10 (very good performance) and 12 (excellent performance). The seven-point grading system was followed only since 2007. Before 2007, a ten-point grading system was followed. The grades included in our study, from years 2002–2006, were converted to seven point grades using the conversion table provided by the Danish Ministry of Education (https://ufm.dk/en/education/the-danish-education-system/grading-system/old-grading-scale). A minimum score of 02 is considered as pass grade. The grade 00 is given if the student has performed extremely poor. Absentees without reason were graded −3. Absentees with an acceptable reason, for example, acute illness, are not graded, but given an opportunity to take the exam in the subsequent year.

We applied a strict sample QC based on the school grades data. We removed individuals who were either younger than 14.5 years or older than 17.5 years at the time of the examinations. The common age group of the students taking the ninth level exams in Denmark is 15 and 16 years. When multiple grades were available for a student, we considered only the grade obtained in the first attempt. We included only individuals who had grades in all the examinations in Danish, English and mathematics chosen for the current study. Also, we removed individuals who had grades in different subjects from different years to avoid heterogeneity that may arise if the student was taught by different teachers, or was at different schools or had different peers between the years.

### School grades in the TEDS cohort

The school grades in the TEDS cohort were from the GCSE exams given at the end of compulsory schooling in UK.^32,33^ The grades were available in three compulsory subjects namely English, mathematics and science. Unlike iPSYCH data, under each subject only one grade was available. The GCSE grades were self-reported by either the participants or their parents. A validation of the self-reported school grades in TEDS has been performed previously by correlating with the grades extracted from national pupil database (NPD) (https://data.gov.uk/dataset/9e0a13ef-64c4-4541-a97a-f87cc4032210/nationalpupil-database) for a subset of the individuals, which showed that the self-reported grades correlated highly (>0.95) with NPD grades.^32^ The GCSE grades ranged between four (G; lowest grade) to 11 (A+; highest grade), with four being the lowest pass grade.

### Occupation and educational attainment in the MVP cohort

Information on primary occupation was collected from all MVP participants through a questionnaire. Only the category of occupation was collected. The participants chose one of the 24 categories (Supplementary Table 17) that matched the best with their primary occupation at the time of recruitment. Individuals who chose the category: Arts, Design, Entertainment, Sport and Media were considered as creative professionals and the rest as controls. Highest level of education achieved by the participants was also collected through questionnaire. The participants chose one of the following answers: less than high school; high school diploma/GED; some college credit, but no degree; associates degree (e.g., AA, AS); bachelors degree (e.g., BA, BS); masters degree (e.g., MA, MS, MBA); professional or doctoral degree. The categories are then converted to number of years of education following International Standard Classification of Education (ISCED) 1997 guidelines (Supplementary Table 18).

### Genotyping and Imputation in the iPSYCH cohort

The source of DNA for genotyping was dried blood spot—two punches of diameter 3.2mm equivalent to a volume of 6 microliter of blood.^24^ The blood spots of the iPSYCH participants were taken from the Danish neonatal screening biobank, which stores blood spot, taken 4-7 days after birth from the heel of the neonate, for all individuals born in Denmark since 1981.^54^ The blood spots were matched with the individuals using the unique identification number that is used across all the registers in Denmark. The extracted DNA was then whole genome amplified in triplicates before genotyping.^55^ The genotyping was performed using Illumina Infinium PsychChip v1.0 array. Genotypes of the markers in the array were identified using well established variant calling pipelines that uses Gencall and Birdseed algorithms.^24^ Following standard QC of the genotyped markers (e.g. call rate > 0.98, MAF>0.01, Hardy-Weinberg equilibrium P value >1×10^−6^), phasing and imputation was carried out. Phasing was performed using SHAPEIT3^56^ and imputation was performed using IMPUTE2^57^ with 1000 genomes phase-3^58^ as the reference panel.

### Genotyping and imputation in the TEDS cohort

Source of DNA for genotyping in the TEDS cohort was saliva collected at the time of recruitment.^32^ The DNA extracted from the saliva was genotyped using either Illumina HumanOmniExpressExome chip or Affymetrix Gene Chip 6.0. Following standard QC procedures, the genotyped markers are phased using EAGLE-2,^59^ followed by imputation using MACH^60^ with Haplotype reference consortium^61^ (release 1.1) as the reference panel. Both phasing and imputation were performed through the Sanger imputation services.^61^ Imputation was performed separately for individuals genotyped using Illumina and those genotyped using Affymetrix. The genotyping chip used was accounted for in the genetic analysis by including a dummy variable for the two chips as covariates.

### Genotyping and Imputation in the MVP cohort

Source of DNA for genotyping in the MVP cohort was peripheral venous blood collected at the time of recruitment or during the follow up visits.^40^ A genotyping chip, called MVP chip (modified Affymetrix Axiom biobank array), was specifically designed for the MVP biobank. The MVP chip contains 723K markers enriched for exome SNPs, validated tag SNPs for diseases including psychiatric disorders and also variants specific to African American and Hispanic populations.^40^ Prephasing was performed using EAGLE v2^62^ and genotypes were imputed using Minimac v3 with the 1000 Genomes Project phase 3, version 5 reference panel.^58^

### Relatedness and population stratification in the iPSYCH cohort

All the individuals involved in the current study were unrelated and had European ancestries. Related pairs of individuals were identified using identity by descent (IBD) analysis using Plink v1.90.^63^ One of each related pair (PIHAT > 0.20) was randomly excluded. PCA was performed in the unrelated individuals using approximately 23,000 imputed variants of high quality (imputation info score > 0.90, MAF>0.05, missing rate < 0.01 and LD independent [r2<0.1]). Among the study individuals, a subset, whose parents and, paternal and maternal grandparents born in Denmark, were identified based on the Danish civil register. These individuals were used as reference to identify population outliers. The first five PCs of the reference individuals were used to construct a five-dimensional ellipsoid with a diameter of eight standard deviations (calculated from the PCs). Those individuals who lied outside the ellipsoid were considered non-Europeans and excluded from the study.

### Relatedness and population stratification in the TEDS cohort

Within each of the genotyped twins, one individual was randomly selected for the current study. Relatedness were estimated among those selected using IBD analysis using Plinkv1.90^63^ and one of each related pair (PIHAT>0.125) were further removed randomly. PCA analysis was performed using EIGENSTRAT^64^ for the unrelated individuals after merging with the 1000 genomes EUR samples.^58^ With the 1000 genomes EUR sample as reference, ancestry outliers were removed iteratively based on the first 10 PCs.

### Relatedness and population stratification in the MVP cohort

Relatedness between MVP participants was inferred using kinship coefficient calculated using software KING.^65^ Related individuals are removed using a kinship coefficient cut off >= 0.0884. Individuals of European ancestries were identified using a machine learning algorithm^66^ that uses information about both self-reported ethnicity as well as principal components derived from PCA of genetic markers. The PCA was performed using EIGENSOFT v.6.^65^

### PCA of school grades

PCA of school grades in the iPSYCH cohort was performed in R using the principal function from the psych R-package. The datasets from years 1990-2006 and 2007-2016 were analyzed separately since the math grade types differed between the two. Both the PCAs yielded six PCs that explained 100% of the variance in the school grades. The PCs were rotated using simplimax algorithm^67^ and then used for the analysis. PCA of school grades in the TEDS cohort was performed in R using prcomp function from the base R-package.

### GWAS

The GWASs in the iPSYCH cohort were performed in Plink v.1.90^63^ using linear regression adjusted for age (age at the time of examination), sex, first 10 PCs, genotyping batches, group variable for PCA of school grades and psychiatric diagnoses. Totally 6,391,200 variants with MAF>0.01 and INFO>0.80 were included for the final analysis.

The GWASs in the TEDS cohort were performed in Plink v.1.90^63^ using linear regression adjusted for age, sex, first 10 PCs and genotyping chips. Totally 5,266,884 variants with MAF>0.01 and INFO>0.80 were included for the final analysis.

### Phenome-wide association analysis

The Phenome-wide association analysis was performed using GWAS atlas,^68^ an online database of GWAS summary statistics. The GWAS atlas database holds summary statistics for 4,571 GWASs (number represent unique studies, but not unique phenotypes). Using variant identifier (rsid) of the index variants in the seven genome-wide loci, we queried the GWAS atlas and obtained all the associations with P value < 0.05. The phenotypes are provided along with category labels such as cognitive and psychiatric. All the associations under cognitive category are provided in the Supplementary Table 4 (educational attainment, though categorized as environmental, is also included in the list).

### SNP-based heritability

The SNP based heritability of the school grades and the E-factors was measured using genome-based restricted maximum likelihood (GREML) analysis implemented in the genome-wide complex trait analysis (GCTA) software.^69^ A genetic relationship matrix (GRM) was constructed using around seven million genetic variants (MAF>0.01; INFO>0.2; Missing rate<0.95) for the whole iPSYCH cohort. The SNP-based heritability was then calculated for only the individuals included in this study (N=30,982) using the GCTA-GREML analysis adjusted for the same covariates as the main GWAS.

### Polygenic scores derivation

In the iPSYCH cohort, polygenic scores for SCZ,^10^ BD,^8^ MDD,^11^ AN,^9^ educational attainment^12^ and intelligence^13^ were derived using effect sizes from summary statistics of the most recently published studies with the largest sample sizes. The polygenic scores for ASD^6^ and ADHD^7^ were derived using effect sizes from summary statistics that were internally generated using a leave-one-out approach. The approach is described in the respective GWASs. Briefly, we divided the sample in to five groups and then performed five GWASs leaving one group away at a time. Each of the GWASs acted as training sample to derive polygenic score in the left-out group. Before calculating polygenic scores, all the summary statistics were similarly processed. The summary statistics were LD clumped using 1000 genome EUR samples^58^ as reference to identify LD-independent variants. Insertion and deletion variants, and variants with ambiguous alleles (A/T, G/C) were removed prior to clumping. The clumped summary statistics were then used for constructing polygenic scores using Plink v1.90.^63^ Ten P-value thresholds were used yielding ten polygenic scores for each trait.

In the TEDS cohort, polygenic scores were constructed for six psychiatric disorders using the same summary statistics that were used for the polygenic score construction in the iPSYCH cohort. In addition, polygenic scores were constructed for E-factors using effect sizes from the GWASs of E-factors in iPSYCH. The summary statistics were LD clumped using 1000 genomes EUR samples as reference to identify LD-independent variants. The clumped summary statistics were then used for constructing polygenic scores using PRSice v2.2.3.^70^

In the MVP cohort, polygenic scores were constructed for E-factors (E1, E2, E3 and E4) using effect sizes from the iPSYCH GWAS of E-factors. The summary statistics were processed using PRS-CS software^71^ to generate weights (posterior SNP effect sizes). Default settings were used for calculating weights using PRS-CS (*γ*-*γ*prior=1; parameter b in *γ*-*γ* prior=0.5; MCMC iterations=1000; number of burn-in iterations=500; thinning of the Markov chain factor=5). Then, based on the derived weights individual level polygenic scores for E-factors were calculated using Plink v2.0^72^ software.

### Polygenic scores analysis

The polygenic score associations were tested using either linear (if analyzing school grades) or logistic regression (if analyzing occupation category). The covariates used in the iPSYCH and TEDS cohorts were same as the ones used in the corresponding GWASs. In the MVP cohort, the following covariates were used: age at recruitment, sex, first 20 ancestral PCs, genotyping batches and number of years of education. The variance explained by the polygenic scores were interpreted using R^2^ (if continuous outcomes: E-factors) or Nagelkerke pseudo-R^2^ (if binary outcomes: occupation). Two regression models were constructed: one with the polygenic score and all covariates included (model 1) and the other with only the covariates included (model 2). The reported R^2^ values were calculated by subtracting the R^2^ of model 2 from R^2^ of model 1.

### Genetic correlations

The genetic correlations were calculated using either GCTA bivariate REML^69^ or LD score bivariate regression.^30^ The genetic correlations between individual school grades, between the E-factors from dataset 1 and E-factors from dataset 2, and pairwise genetic correlations between the E-factors (after combing both the datasets) were all calculated using GCTA using individual genotypes. The genetic correlations between the E-factors and other traits (years of education, intelligence and the six psychiatric disorders), and between the iPSYCH E-factors and TEDS E-factors were calculated using LD score regression using GWAS summary statistics. We used the precalculated LD scores available from the LD score regression website. All the summary statistics were processed in a similar way. Only the HapMap variants (SNP list available from LD score regression website) were used for the analysis. The reference and effect alleles in the summary statistics are aligned with that of HapMap list. Then, the summary statistics files were munged using the munge script from LDSC software (with default settings). The munged files were then used to estimate genetic correlations.

We tested if the genetic correlation of E1 with educational attainment is significantly different from the genetic correlation of E1 with intelligence by block jackknife method implemented in LD score regression.^73^ In brief, LD score regression generates standard errors for heritability and for genetic correlation via a jack-knife bootstrap by repeatedly estimating the statistic of interest while excluding different blocks of data each time. The same approach can be used to generate robust standard errors for the difference between two genetic correlations, enabling robust estimation of the significance of this difference.

### Multiple testing corrections

The statistical significance in all the analyses was assessed after multiple testing correction. The P value threshold in each of the analyses was decided based on the number of unique hypotheses tested. In the analysis of the heritability of E-factors (N=4), the P value threshold was set to 0.01 (0.05/4). In the GWAS analysis, the P value threshold was set to 1.25×10^−8^ (5×10^−8^ / 4). In the analysis of genetic correlations and polygenic score associations of E-factors (N=4) with educational attainment and intelligence (N=2), the P value threshold was set to 0.006 (0.05/8). In the analysis of phenotypic associations, genetic correlations and polygenic score associations between the E-factors (N=4) and the psychiatric disorders (N=6), the P value threshold was set to 0.002 (0.05 /24). In the analysis of phenotypic associations and polygenic score associations of math and language (N=2) with psychiatric disorders (N=6), the P value threshold was set to 0.004 (0.05/12). In the analysis of genetic correlations and polygenic score associations of iPSYCH E-factors (N=4) with TEDS E-factors (N=2), the P value threshold was set to 0.006 (0.05/8). In the analysis of polygenic score associations of TEDS E-factors (N=2) with psychiatric disorders (N=6), the P value threshold was set to 0.004 (0.05/12). In the analysis of polygenic score associations of E2 (N=1) with occupations (N=24), the P value threshold was set to 0.002 (0.05/24).

## Notes

### Summary of Updates

One of the author names was missed during the first upload. It's now corrected.

