## Supplementary Information for "Genome-wide association study of school grades identifies a genetic overlap between language ability, psychopathology and creativity"

**Supplementary Information** **1**

**Bibliography** **14**

### 1.1 Supplementary Figures

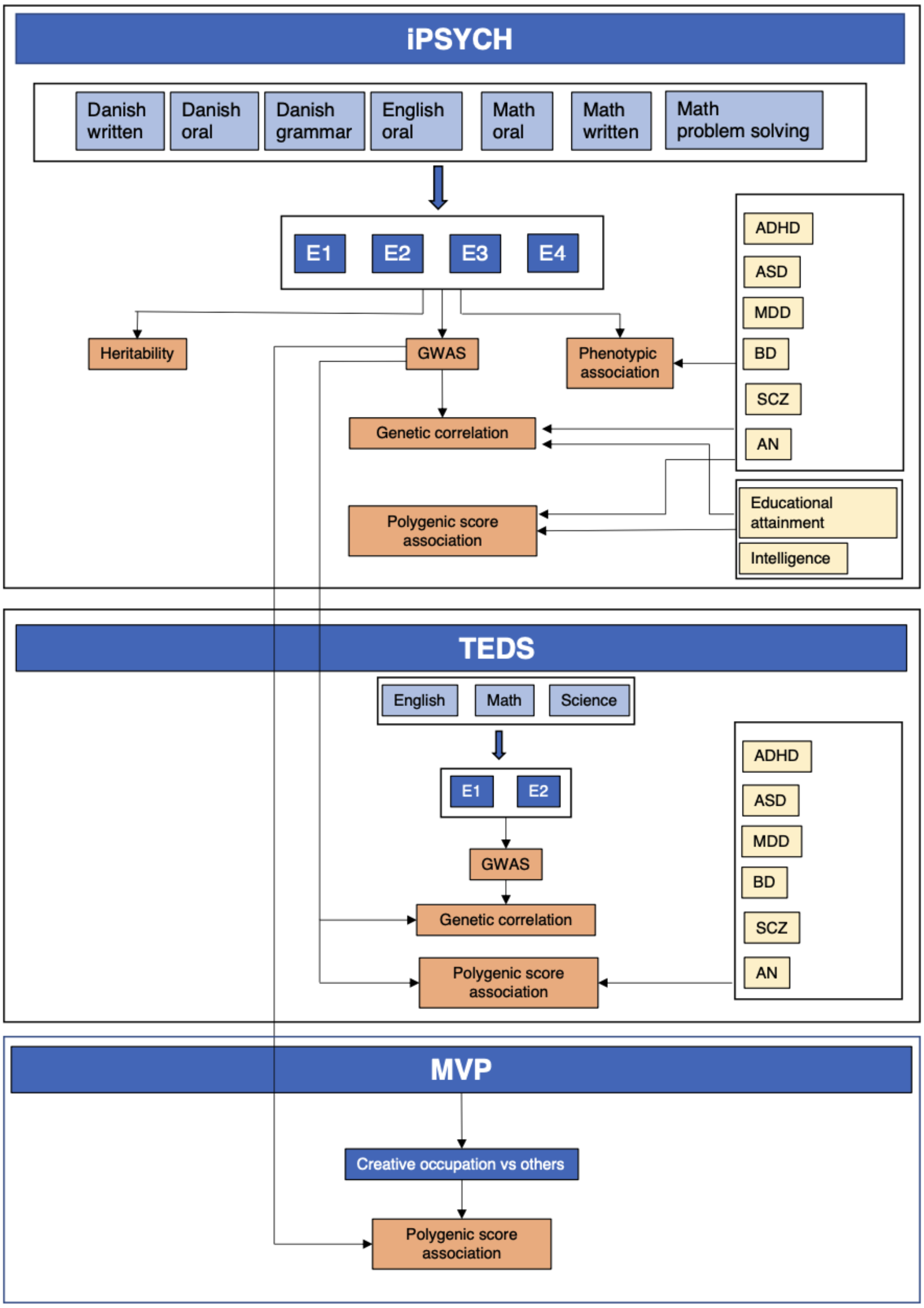

**Supplementary Fig. 1 | Overall study design**

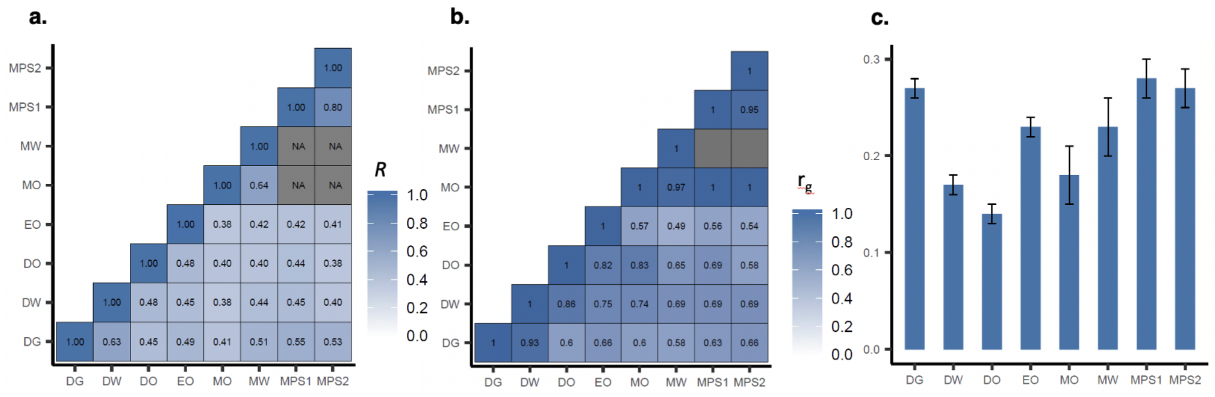

**Supplementary Fig. 2 | Phenotypic and genetic correlations and SNP-based heritability of subject specific grades.** **a.** Phenotypic correlations across school grades. Pearsons correlation coeffients are plotted. NA values correspond to non-overlapping sets **b.** Genetic correlation matrix of school grades. $r_g$ values, calculated using bivariate GREML analysis in GCTA, are plotted; **c.** SNP heritability of school grades, calculated using GREML analysis in GCTA, are plotted; DG=Danish grammar; DW= Danish wriiten; DO=Danish oral; EO=English oral; MO=Math oral; MW=Math written; MPS1= Math problem solving1; MPS2=Math problem solving 2

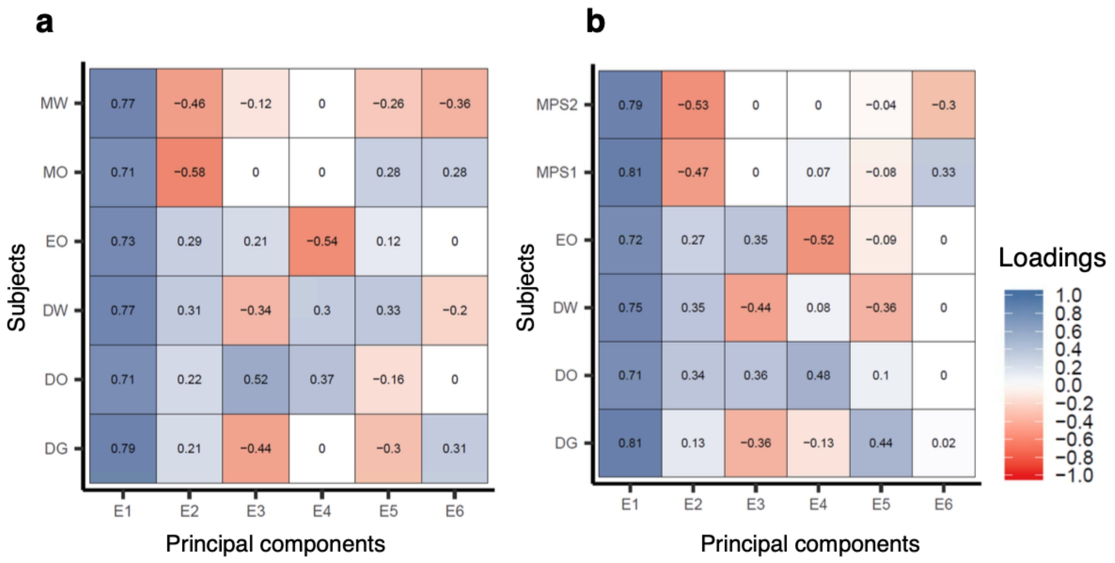

**Supplementary Fig. 3 | PC loadings in controls.** The figures here are same as the main figures 1a and 1b, except that here the PCA was performed only in the controls sample (N=12,487). This is done to ensure that the loadings structure is not affected by including individuals with psychiatric disorders in the PCA analysis. **a.** PC loadings of the E-factors in dataset 1 [2002-2006; N=3,816]. **b.** PC loadings of the E-factors in dataset 2 [2007-2016; N=8,671]; DG=Danish grammar; DO=Danish oral; DW= Danish written; EO=English oral; MO=Math oral; MW=Math written; MPS1= Math problem solving 1 ; MPS2=Math problem solving 2

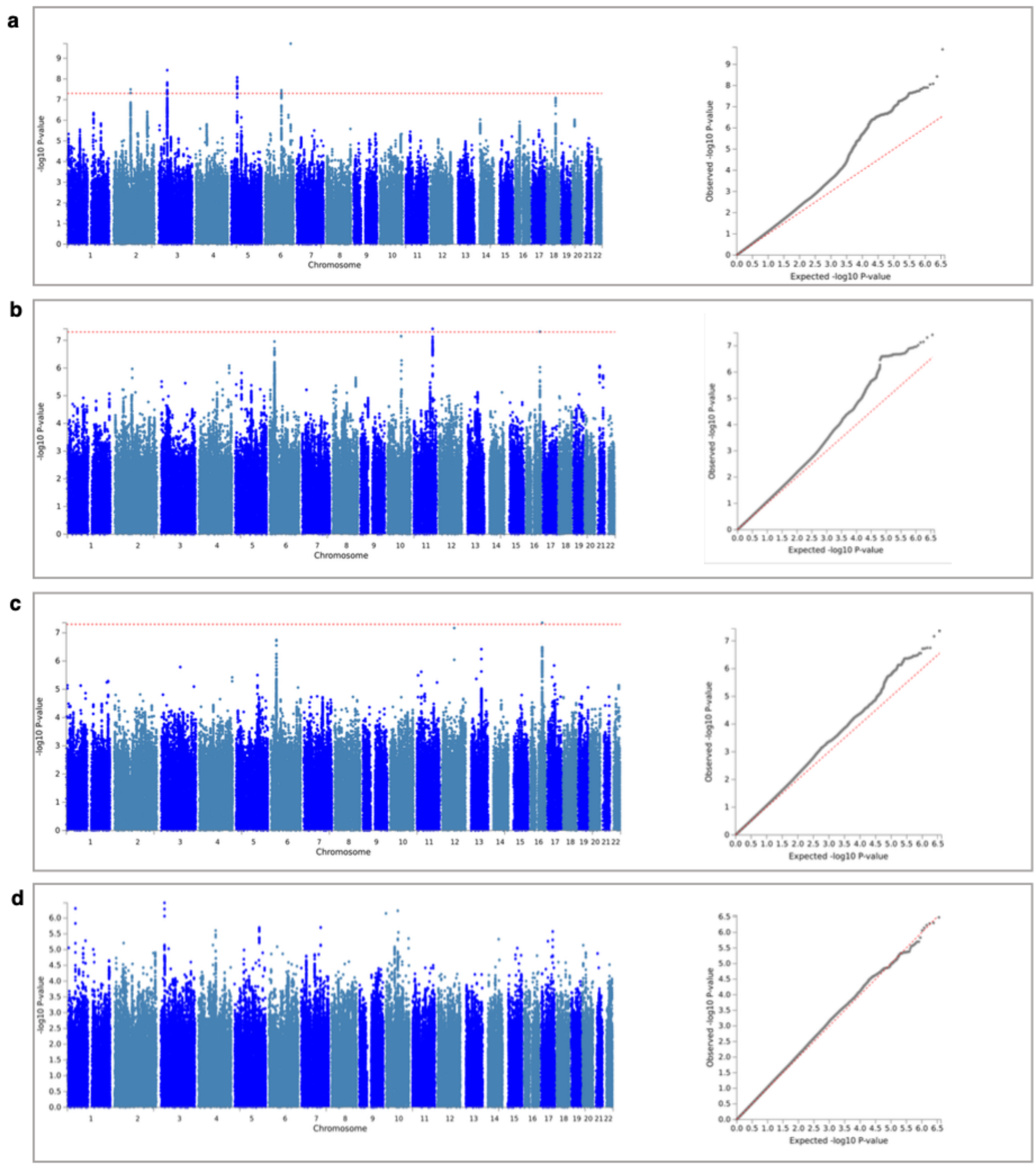

**Supplementary Fig. 4** | **Manhattan and QQ plots** of the GWASs of E1 (a), E2 (b), E3 (c), and E4 (d). The plots are created using online tool: FUMA (fuma.ctglab.nl). The horizontal dotted red lines in the Manhattan plots correspond to genome wide significant threshold (P< $5x10^{-8}$). The diagonal dotted red lines in the QQ plots correspond to expected correlation when the observed results have a null distribution.

### 1.2 PCA of school grades in the iPSYCH cohort

To decompose the subject specific school grades in to distinct factors, we performed a principal component analysis (PCA) of the school grades. In the first dataset (i.e. grades from exams conducted between 2002 and 2006), for each individual we had totally six different grades namely Danish written, oral and grammar, English oral, mathematics written and oral grades. Totally 11,284 individuals from the first dataset were included after QC. PCA yielded six principal components (PCs) that together explained 100% of the variance (Supplementary Table 1).

In the second dataset (i.e. grades from exams conducted between 2007 and 2016), for each individual we had totally six different grades namely Danish written, oral and grammar, English oral, mathematics problem solving 1 (without any digital assistance) and problem solving 2 (with digital assistance such as calculator). Totally 19,698 individuals from the second dataset were included after QC. Since the math grades are not the same in the first and second datasets, we performed PCA separately. The PCA yielded six PCs that together explained 100% of the variance in the school grades (Supplementary Table 1).

The subject loadings of the first four PCs are comparable between the two datasets. Moreover, we found that the first four corresponding PCs of the two datasets had near perfect genetic correlations (rg $\sim$ 1), calculated using GCTA bivariate GREML analysis.[1] Since the fourth and fifth PCs in the two datasets explained relatively less variance and did not align between the two datasets, we did not analyze them further and restricted our further analyses to only the first four PCs in the two datasets.

We call the first four PCs as E-factors (Education factors) to differentiate them from the ancestral PCs that are used to correct for population stratification. The factor E1 in both the datasets explained the largest amounts of variance (as one would expect in a PCA) and correlated almost similarly with all the individual subject grades (Fig. 1b, 1b). Hence, E1 measured the overall school performance. A higher E1 score in an individual indicates that the individual performed equally well in all the six grades. E1 is same as the mean across all the grades (Pearson's correlation, $R$, between E1 scores and mean grades was 0.998 and 0.996 in the first and second datasets respectively).

The subject loadings of the factor E2 in the two datasets were intriguing. The factor E2 correlated positively with Danish (oral) and English (oral, written and grammar) grades and negatively with mathematics grades (written and oral in dataset1 and problem solving 1 and 2 in dataset2; Fig 1a, Fig 1b). Hence, the factor E2 separated individuals based on their differential performance in language and math. Those with higher E2 scores had better language grades relative to their math grades and those with lower E2 scores had the opposite. It is important to note that E2 is a relative measure. An individual with a poor grade in both language and math can still have a higher E2 score if the individuals language grades are relatively better than their math grades (i.e. the individual is poor in both language and math, but slightly better in language compared to their math). This was the case in many of the individuals with ADHD.

The factor E3 correlated mostly with only Danish and English grades, but not with mathematics grades. The loadings were opposite in directions between written and oral grades. E3 correlated positively with Danish oral and English oral grades and negatively with Danish written and grammar grades (Fig 1a, 1b). Hence, E3 separated individuals based on their differential performance in written and oral language (Danish and English) grades. Those with higher E3 score had better oral grades relative to their written grades and those with lower E3 scores had the opposite. Similar to E2, E3 is also a relative measure.

Similar to E3, the factor E4 also correlated only with language (Danish and English) grades, but not with mathematics grades, in particular the factor correlated with only oral grades, but not written grades (minimal correlation with Danish written and grammar). The loadings were opposite between Danish and English oral grades (Fig 1a, Fig 1b). E4 correlated positively with Danish oral and negatively with English oral grades. Hence, E4 separated individuals based on their differential performance in Danish and English oral grades, possibly reflecting a difference between native and foreign language speaking skills. Those with higher E4 scores had better Danish oral grades relative to their English oral grades and those with lower E4 scores had the opposite. Similar to E2 and E3, E4 is also a relative measure.

### 1.3 Association of sex with E-factors

All the four E-factors were significantly associated with sex (Supplementary Table 2). E1 was associated negatively with sex, which suggested that females performed significantly better than males in the overall performance (Beta=-0.13; SE=0.01; P=3x10$^{-31}$; females were coded as reference). E2 was associated negatively with sex, which suggested that, when comparing between math and language performances, females scored better in language than in math whereas males scored better in math than in language (Beta= -0.56; SE=0.01; P<1x10$^{-300}$). E3 was associated positively with sex, which suggested that, when comparing between written and oral exams, females performed better in written than in oral whereas males performed better in oral than in written exams (Beta=0.29; SE=0.01; P=2.7x10$^{-125}$). E4 was associated negatively with sex, which suggested that, when comparing between English and Danish exams, females performed better in English than in Danish whereas males performed better in English than in Danish exams (Beta=-0.30; SE=0.01; P=1.6x10$^{-136}$). Overall, the associations of E2, E3, E4 were stronger than that of E1 suggesting that subject specific grades, compared to overall grades, show more heterogeneity with regard to sex. The strongest association of all was observed for E2. The effect size of E2 (Beta=-0.56) is four times the effect size of E1 (Beta=-0.13) suggesting that sex-based differences were strongest with regard to math and language grades.

We further looked into the math and language specific associations with sex only in the controls similar to the analyses described for psychiatric disorders. The results showed that the males scored better in math compared to females (Beta=0.19; SE=0.01; P=8.1x10$^{-63}$) and females scored better in language compared to males (Beta=-0.31; SE=0.01; P=3.4x10$^{-158}$). Notably, the language effect size is larger than math effect size, which might explain why overall performance is higher in females despite that females scored lower in math compared to males.

### 1.4 Association of age with E-factors

All the four E-factors were significantly associated with exam age (calculated at the time of the examinations; Supplementary Table 2). The age of the individuals included in the study ranged between 14.5 years to 17.5 years. E1 was associated negatively with age suggesting that individuals who performed better in the exit exam were significantly younger compared to individuals who performed poorer in the exit exam (Beta=-0.33; SE=0.01; P=1.1x10$^{-141}$). The results only indicate that poor performers finished compulsory schooling later compared to better performers. The results do not indicate any association between brain age and school performance.

E2 was associated positively with age suggesting that younger individuals (i.e. individuals who finished compulsory school early) scored better in math relative to language whereas older individuals (i.e. individuals who finished compulsory schooling late) scored better in language relative to math (Beta=0.07; SE=0.01; P=3x$10^{-8}$).

E3 was associated positively with age suggesting that younger individuals scored better in oral exam relative to written whereas older individuals scored better in written exams relative to oral (Beta=0.07; SE=0.01; P=4.4x$10^{-8}$)

E4 was associated positively with age suggesting that younger individuals attending exit exams scored better in English relative to Danish whereas older individuals attending exit exams scored better in Danish relative to English (Beta=0.08; SE=0.01; P=1.2x$10^{-9}$)

Though all E-factors were significantly associated with exam age, the association was stronger only for E1. The effect sizes of E2, E3 and E4 were all minimal (Beta <0.10) whereas the effect size of E1 is -0.35. Hence, age factor influences only the overall performance, but not the subject specific performances substantially.

Also, we tested the association of age with math and language grades only in the controls. Both math and language grades were negatively associated with age and the effect sizes were almost similar with math effect size (Beta=-0.33; SE=0.01; P=1.1x$10^{-141}$) being slightly higher than language effect size (Beta=-0.30; SE=0.01; P=3.7x$10^{-123}$). Hence, individuals who finished compulsory schooling earlier scored higher in both math and language compared to individuals who finished compulsory schooling later.

### 1.5 Pleiotropic associations and annotations of the risk variants in the genome-wide loci

The sample size of our GWASs, compared to previous GWASs of educational attainment[2] and intelligence,[3] were small by multiple orders of magnitude. Importantly, gene discovery was not the main aim of our study. However, we discuss here few of the interesting genes mapped to the seven genome-wide loci we identified. We queried the haploreg database (v4.1)[4] using the index SNPs at the seven loci and explored the mapped genes for their relevance to brain functions. Also, we discuss some of the interesting pleiotropic associations of the index variants that were identified through phenome-wide association analysis (Methods).

#### Genome-wide loci identified for E1

##### Locus 2q11.2

1. **Pleiotropic associations**: The index variant, rs11895772, in this locus is strongly associated with years of schooling (P=2.9x$10^{-24}$). The variant is also significantly (P<0.05) associated with 21 other cognitive phenotypes including cognitive performance (P=7x10-14), verbal and numerical reasoning (P=7.2x$10^{-9}$), fluid intelligence (P=5.2x$10^{-7}$) etc. (Supplementary Table 4a). Also, the variant is also strongly associated with multiple metabolic phenotypes such as body mass index, waist hip ratio, body fat percentage etc.

2. **Annotations**: The variant rs11895772 is located in a highly regulatory region: 5' UTR of gene *LONRF2* and is in LD (r2>0.80) with numerous variants. The locus overlaps with enhancers and promoters specific to multiple tissues including brain and also contains multiple transcription factor binding motifs. Among the LD variants, five lie in exonic regions, of which two are missense and three synonymous.

Both the missense variants lie in the exon of *LONRF2*. Among the three synonymous variants, two lies in the exon of *LONRF2* and one in the exon of *CHST10*. Also, many of the LD variants are located in the intronic regions of *LONRF2* and *CHST10*. As the region spanning the locus is highly regulatory in nature, all the LD variants are eQTLs for multiple genes including *LONRF2* and *CHST10* in multiple tissues including brain tissues.

3. **Gene functions**: Not many genes are located in the vicinity of the locus 2q11.2. Among those present, the genes *LONRF2*, *CHST10* and *NMS* are brain-related. The gene *CHST10* codes for a sulfotransferase enzyme protein called carbohydrate sulfotransferase 10. The enzyme act on human natural killer-1 (HNK-1) glycan, which is involved in neurodevelopment and synaptic plasticity.[5] The gene *LONRF2* codes for protein neuroblastoma apoptosis-related protease whose expression is highly brain-specific.[6] Its functional significance is unknown. The gene *NMS* codes for a neuropeptide protein called neuromedin S. The neuropeptide is involved in regulation of circadian rhythm and, has anorexigenic and antidiuretic actions.[7]

**Locus 3p21.31**

1. **Pleiotropic associations**: The index variant, rs7613360, in this locus is strongly associated with years of schooling (P=$1.4x10^{-52}$). The variant is also significantly associated with 27 other cognitive phenotypes (Supplementary Table 6). Also, the variant is strongly associated with various other phenotypes under domains such as metabolic, psychiatric, reproductive, immunological and addiction phenotypes. Among them, the associations with BMI (P=$2.9x10^{-28}$), reticulocyte count ($2.5x10^{-14}$), age at first live birth (P=$6.8x10^{-16}$), age at menarche (P=$5.8x10^{-15}$), age first had sexual intercourse (P=$4.9x10^{-14}$), chronotype (P=$4.4x10^{-11}$), miserableness (P=$1.9x10^{-8}$) and morning person (P=$5x10^{-10}$) are notable.

2. **Annotations**: The variant, rs7613360, is in LD (r2>0.80) with 17 other variants. Among them, two are synonymous exonic variants located in genes *CAMKV* and *CTD2330K9.3* respectively; 11 are intronic and 5 are non-coding variants. The locus is strongly regulatory in nature as it overlaps with multiple promoters and enhancers specific to multiple tissues including brain and also contains multiple transcription factors binding motif. Consequently, all the 18 variants are strong eQTLs for multiple genes including *CAMKV*, *MST1R* and *MON1A* in multiple tissues including brain tissues.

3. **Gene functions**: Multiple genes are located in the locus 3p21.31. Notable among them are the genes *CAMKV*, *SEMA3F* and *SEMA3B*. The gene *CAMKV* encodes a synaptic protein called calmodulin kinase-like vesicle-associated, which regulates synaptic transmission and plasticity thereby regulating learning and memory processes.[8] The gene *SEMA3F* encodes semaphorin 3F, which is signaling protein involved in axon guidance during neuronal development.[9] The gene *SEMA3B* encodes semaphorin 3B, which is a signaling protein involved in axonal guidance during brain development, particularly is critical for anterior commissure development.[10]

**Locus 5p13.2**

1. **Pleiotropic associations**: The index variant, rs696732, is significantly associated with neither educational attainment (P=0.9) nor intelligence (P=0.07). Also, none

of the cognitive phenotypes showed significant associations with rs696732 in the PheWAS analysis. The minor allele frequency of rs696732 is 0.27 in the iPSYCH cohort and 0.29 in the Europeans subset of 1000 genomes. Also, multiple variants in LD with rs696732 also showed strong associations with E1 as can be seen in the regional association plot (Extended dataset 2). Hence, the association with E1 at this locus is unlikely to be a false positive signal, but might represent a novel association driven through mechanisms unique to school grades or through environment unique to the iPSYCH cohort. Interestingly, based on the top PheWAS hits of rs696732, the locus is strongly related to respiratory and immunological phenotypes. For example, the variant is strongly associated with traits: Blood clot, DVT, bronchitis, emphysema, asthma, rhinitis, eczema, allergy diagnosed by doctor: Hay fever, allergic rhinitis or eczema (P=1.1x$10^{-37}$), asthma, hay fever or eczema (P=5.5x$10^{-19}$), Doctor diagnosed hay fever or allergic rhinitis (P=5.1x$10^{-13}$) and eosinophil count (8.3x$10^{-10}$). Based on these pleiotropic associations it might be possible that the locus affects school performance not through brain related mechanisms but by causing recurrent respiratory illness such as hay fever in the school going students (which is common in Denmark[11]). Studies have shown that hay fever affects cognitive performance in school going children.[12]

2. **Annotations**: The variant, rs696732, is an intronic variant located in the gene *CAPSL*. Totally 18 variants are in LD (r2>0.80) with rs696732, among which, 12 are intronic (in the genes *CAPSL* and *UGT3A1*) or non-coding (one variant 3' UTR of *UGT3A1* and the rest located 3' of *RP11-79C6.3*). The locus overlaps with enhancer histone marks notably enhancers specific to lungs, blood and thymus. Multiple transcription factor binding motifs are present in the locus. And all the variants have at least one eQTL association with gene *LMBRD2* in the GTEx lung tissue.

3. **Gene functions** Among the genes located around locus 5p13.2, none seem to have any brain-related functions. However, one of the genes, *IL7R* has immune related function. *IL7R* codes for interleukin-7 receptor alpha chain. *IL-7* receptors are present in the immune system cells such as B-cells and T-cells.[13] The gene seems to the most relevant one at this locus, given the PheWAS associations with immunological phenotypes.

**Locus 6q16.1**

1. **Pleiotropic associations** The index variant, rs2388334, is strongly associated with years of schooling (P=2.1x$10^{-39}$) and intelligence (P=3.6x$10^{-29}$). The variant is also significantly associated with 27 other cognitive phenotypes. Apart from cognitive associations, the variant is also strongly associated with other groups of traits namely metabolic, psychiatric and neurological traits. Among them, the associations with BMI (P=1.4x$10^{-15}$), schizophrenia/bipolar disorder (P=1.3x$10^{-8}$), bipolar disorder (P=4.9x$10^{-8}$), ASD (P=1.0x$10^{-6}$), brain volume: left middle temporal (P=2.1x$10^{-6}$) and brain volume: right insula (P=7.1x$10^{-6}$) are notable.

2. **Annotations**: The variant rs2388334 is a non-coding variant located 5' of *RP11-111D3.2*. The variant is in LD (r2>0.80) with 25 variants, all of which are non-coding located either 3' of *MIR2113* or 5' of *RP11-111D3.2*. The locus overlaps with multiple enhancers and promoters in multiple tissues including brain. The region also contains multiple transcription factor binding motifs. Unlike other loci, no

eQTL associations were found for the variants in the locus 6q16.1 except for a single eQTL association between variant rs9385269 and gene *CCR10* in the peripheral monocytes.

3. **Gene functions**: Only one gene, *MIR2113*, is located in the vicinity of locus 6q16.1. The gene *MIR2113* codes for microRNA 2113. Micro RNAs are involved in post-transcriptional regulation of gene expression. The function of this microRNA has not been studied so far.

### Genome-wide loci identified for E2

#### Locus 11q23.2

1. **Pleiotropic associations**: The index variant, rs4547132, at the locus 11q23.2 is significantly associated with 14 cognitive phenotypes including years of schooling (P=0.03). However, all the associations are only moderately significant (lowest P value is 0.00001 for phenotype: Symbol digit substitution test - Number of symbol digit matches attempted). The variant is, however, strongly associated with various other traits including psychiatric, metabolic and addiction traits. Among them, the associations with ever smoker ($P=1.2x10^{-24}$), first PC of the four risky behaviors ($P=2.6x10^{-13}$), number of sexual partners ($P=1.8x10^{-7}$), neuroticism ($P=1.6x10^{-6}$), depressive symptoms (P=0.0001) and schizophrenia (P=0.004) are notable.

2. **Annotations**: The index variant, rs4547132, is an intronic variant located within gene *RP11-629G13.1.* The variant is in LD (r2>0.6) eight variants, of which six are intronic located within gene *NCAM1* and two are non-coding located 3 of gene *RP11-629G13.1.* The locus overlaps with multiple promoters and enhancers in multiple tissues including brain and placenta. Also, the locus contains multiple transcription factor binding motifs.

3. **Gene functions**: Not many genes are located in this locus. Among those found, the most notable gene is *NCAM1*, which codes for neural cell adhesion molecule 1. The protein is involved in development and differentiation of nervous system.[14]

#### Locus 16q23.3

1. **Pleiotropic associations**: The index variant, rs11150461, is significantly (P<0.05) associated with 5 cognitive phenotypes, though all the associations are only borderline significant. The smallest P value (0.007) was seen for the phenotype, Prospective memory test - Time to answer. The variant is also associated multiple other phenotypes under categories such as metabolic, psychiatric etc. Some of them are: BMI ($P=2.1x10^{-11}$), waist circumference ($P=6.1x10^{-11}$), risk taking (P=0.005) and schizophrenia (P=0.01).

2. **Annotations**: The variant rs11150461 is a non-coding variant located 5' of gene snoU13. The variant is in LD (r2>0.6) with 16 other variants, all of which are non-coding located 5' of snoU13. The region overlaps with few enhancers in tissues blood and adrenal glands, but no overlap with any promoters. The locus has multiple transcription binding motifs. No eQTL associations are reported so far for the variants in this locus.

3. **Gene functions**: Not many genes are located at this locus. Among those present, one gene, *CDH13*, is brain related. The gene *CDH13* encodes for a protein called cadherin 13, which acts as a negative regulator of axon growth during neural differentiation.[15]

#### Genome-wide locus identified for E3

##### Locus 16q22.1

1. **Pleiotropic associations**: The index variant, rs4985376, is significantly (P<0.05) associated with five cognitive phenotypes, though all the associations are only borderline significant. Notably the variant is associated with handedness phenotypes: right handedness (P=0.03), right handedness (P=0.04), handedness (P=0.04). The variant is strongly associated with multiple metabolic, reproductive and respiratory phenotypes such as BMI (P=1.1x$10^{-17}$), age at menarche (P=5.6x$10^{-15}$) and FEV1 (P=1.8x$10^{-11}$).

2. **Annotations**: The variant rs4985376 is an intronic variant located in gene *WWP2*. The variant is in LD (r2>0.8) with 21 other variants, all of which are intronic variants located in gene *WWP2*. The region is regulatory in nature. It overlaps with multiple promoters and enhancers in multiple tissues including brain tissues. Also, the region contains multiple transcription binding motifs. All the variants have multiple eQTL associations with multiple genes.

3. **Gene functions**: Many genes are located in this locus. Among them, the genes *NFAT5* and *EXOSC6* are brain-related. *NFAT5* codes for protein called Nuclear factor of activated T-cells 5, which is a transcription factor and is involved in transcriptional regulation of immune and inflammation related genes. The gene is highly expressed in fetal and adult brain, the fetal brain expression being 10-fold higher than adult brain expression.[16] The gene *EXOSC6* codes for a protein called exosome component 6, which is a subunit of exosome and is involved in mRNA degradation.[17]

### 1.6 Association of E2, E3 and E4 with years of education and intelligence

The genetic correlations of E2, E3 and E4 with educational attainment[2] and intelligence[3] were lower compared to that of E1 (Fig. 2a; Supplementary Table 6). The polygenic score analysis mirrored the genetic correlations (Fig 2b, 2c; Supplementary Table 7). Among the genetic correlations, only the correlations of E2 with educational attainment and intelligence, and E4 with intelligence remained statistically significant after multiple testing correction (P<0.006). All of the polygenic score associations, however, were statistically significant after multiple testing correction.

E2 correlated negatively with both educational attainment ($r_g$ =-0.12; SE=0.03; P=0.002) and intelligence ($r_g$ =-0.19; SE=0.04; P=0.00001) suggesting that the common variants associated with better performance in language relative to math are also associated with lesser number of years of education and poor intelligence. E3 correlated positively with years of education ($r_g$ =0.08; SE=0.03; P=0.008), but negatively with intelligence ($r_g$ =-0.10; SE=0.04; P=0.01) suggesting that the common variants associated with better performance in oral exam relative to written exam are also associated with higher number of years of education, but poor intelligence. E4 correlated negatively with both years of education ($r_g$ =-0.1; SE=0.04; P=0.03) and intelligence ($r_g$ =-0.32; SE=0.05; P=1.4x10-8) suggesting that the common variants associated with better performance in Danish relative to English are also associated with lesser number of years of education and poor intelligence.

### 1.7 Association of E3 and E4 with psychiatric disorders

Unlike E1 and E2, we didnt see statistically significant associations for E3 and E3 with the psychiatric disorders either in the genetic correlation analysis (Fig. 3b; Supplementary Table 9) or in the polygenic score association analysis (Fig. 3c; Supplementary Table 10) except for the genetic correlation between ADHD and E3 ($r_g$ =0.17; SE=0.05; P=0.002). However, we observed few significant associations at the phenotypic level (Fig. 3a; Supplementary Table 8).

We observed significant differences in the E3 scores between cases and controls with regard to MDD, ADHD and ASD; also, in the E4 scores with regard to ADHD and ASD. Individuals with MDD scored had significantly lower E3 scores compared to controls suggesting that individuals with MDD performed better in oral exams relative to written in comparison to controls (Beta=-0.04; SE=0.01; P=0.0008). Individuals with ADHD and ASD both had significantly higher E3 scores compared to controls suggesting that they performed significantly better in oral exams relative to written in comparison to controls (ADHD: Beta=0.08; SE=0.01; P=$2.6\text{x}10^{-7}$; ASD: Beta=0.12; SE=0.01; P=$8.6\text{x}10^{-13}$).

We observed significant differences in E4 scores between cases and controls with regard to ADHD and ASD. Individuals with ADHD and ASD had significantly lower E4 scores compared to controls suggesting that they performed poorer in Danish relative to English in comparison to controls (ADHD: Beta=-0.08; SE=0.01; P=$1.1\text{x}10^{-7}$; ASD: Beta=-0.18; SE=0.01; P=$1.2\text{x}10^{-25}$).

We observed a significant positive genetic correlation between ADHD and E3 suggesting that common variants associated with better performance in oral exams relative to written exams are also associated with increased risk for ADHD (rg=0.17; SE=0.05, P=0.002).
