## Supplementary Dataset 1 for "Genome-wide association study of school grades identifies a genetic overlap between language ability, psychopathology and creativity"

#### Supplementary dataset 1 l Regional association plots

The regional association plots of the seven genome-wide loci (four for E1, two for E2 and one for E3) are in this file. The plots are made using online tool, LocusZoom (locuszoom.org). Genotypes of 1000g Phase 3 European subset were used for LD calculation.

In all the plots, in the X axis basepair positions (hg19), in the left Y axis $-\log_{10}$ P values and in the right Y axis recombination rates are plotted. The index variant (i.e. the most significant variant) is shown as a diamond and is labelled. The points are colored according to their LD ($r^2$) with the index variant. The genes located within the regions are displayed as gene tracks (thick line correspond to exon and thin line, intron). The arrowed end correspond to the 3' end of the gene.

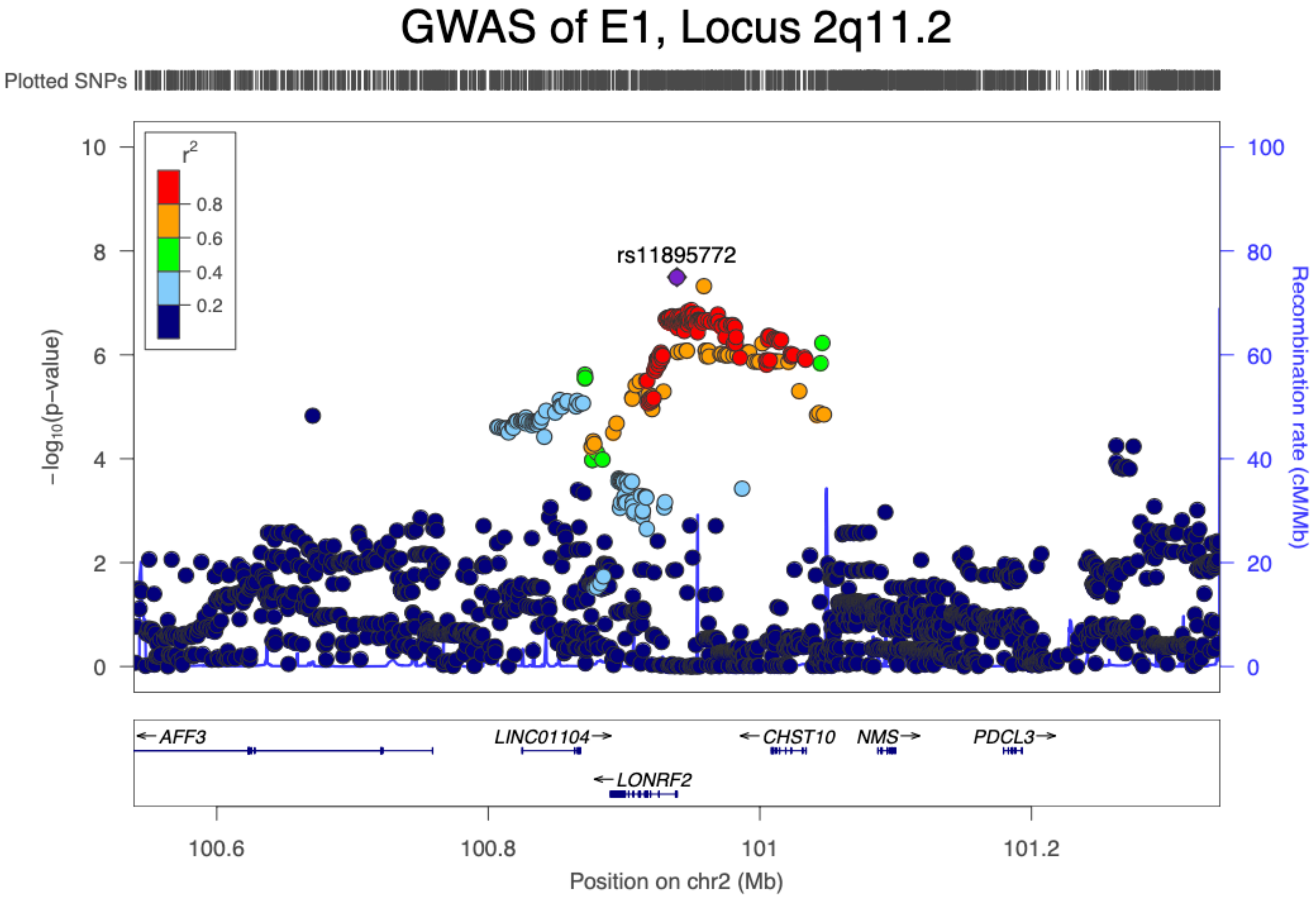
GWAS of E1, Locus 2q11.2
Plotted SNPs
r2
0.8
0.6
0.4
0.2
rs11895772
−log10(p−value)
Recombination rate (cM/Mb)
0
2
4
6
8
10
0
20
40
60
80
100
AFF3
LINC01104
CHST10
NMS
PDCL3
LONRF2
100.6
100.8
101
101.2
Position on chr2 (Mb)

### GWAS of E1, Locus 3p21.31

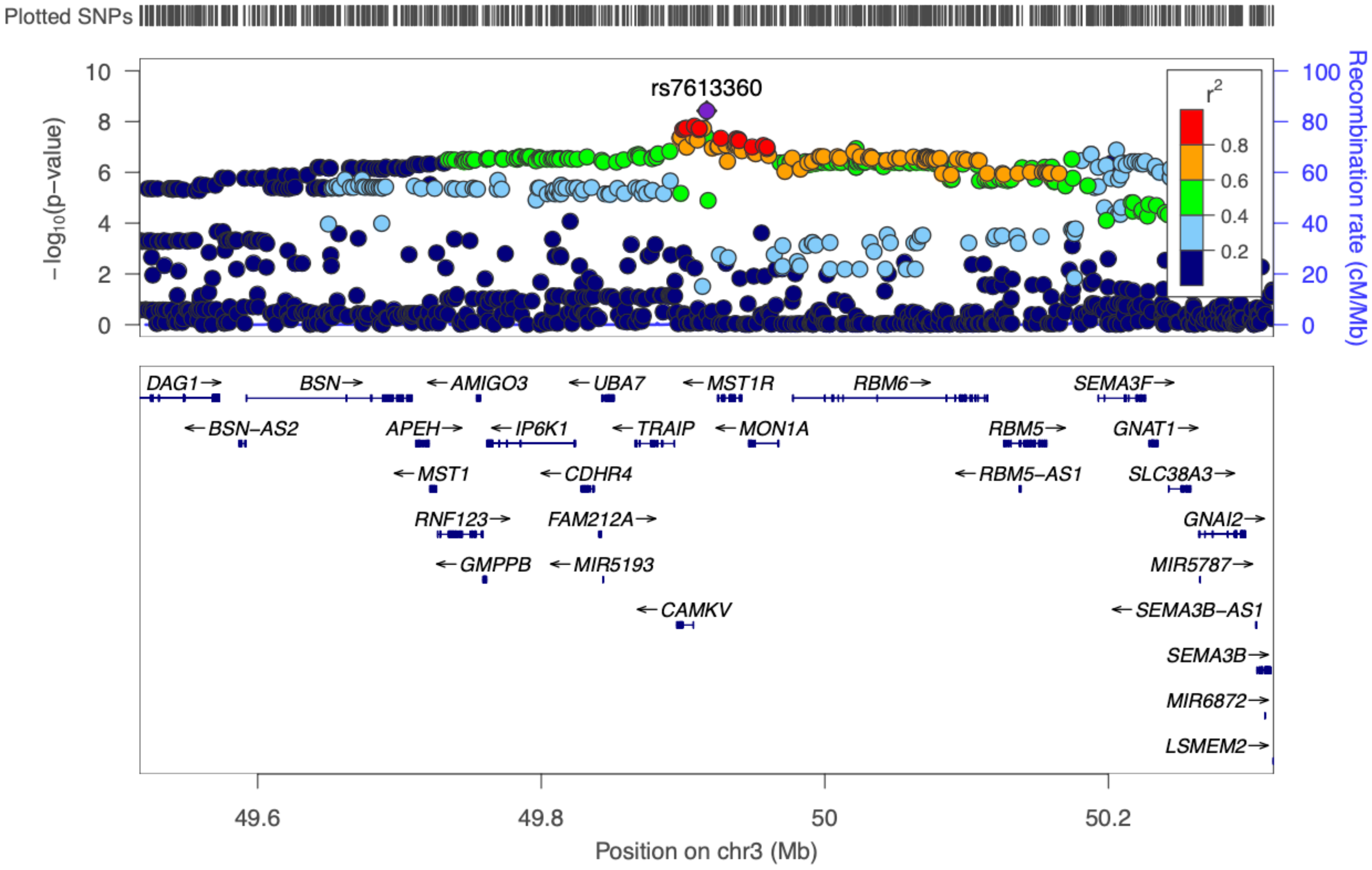

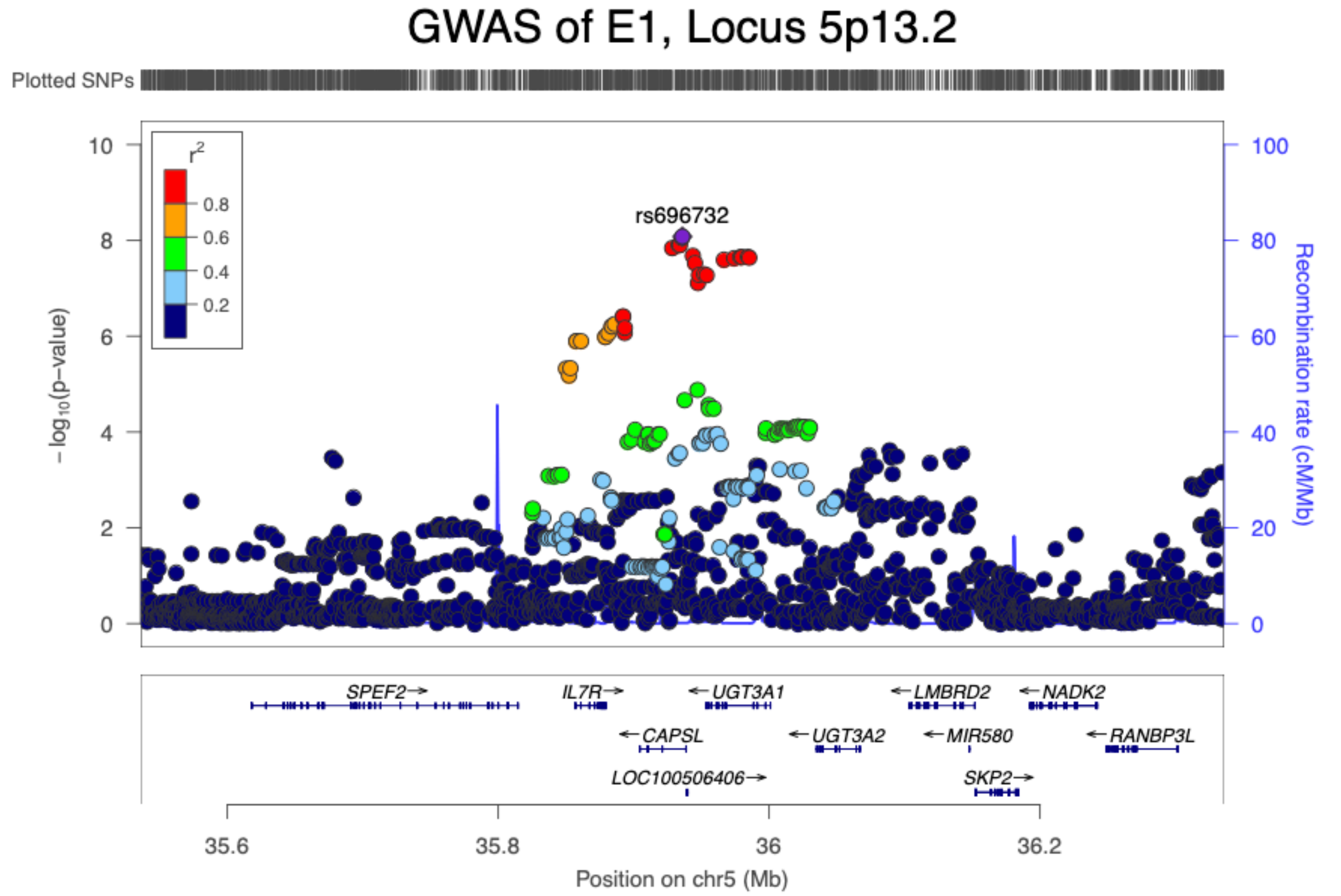
GWAS of E1, Locus 5p13.2
Plotted SNPs
rs696732
r²
0.8
0.6
0.4
0.2
10
8
6
4
2
0
−log10(p−value)
100
80
60
40
20
0
Recombination rate (cM/Mb)
SPEF2→
IL7R→
←UGT3A1
←LMBRD2
←NADK2
←CAPSL
←UGT3A2
←MIR580
←RANBP3L
LOC100506406→
SKP2→
35.6
35.8
36
36.2
Position on chr5 (Mb)

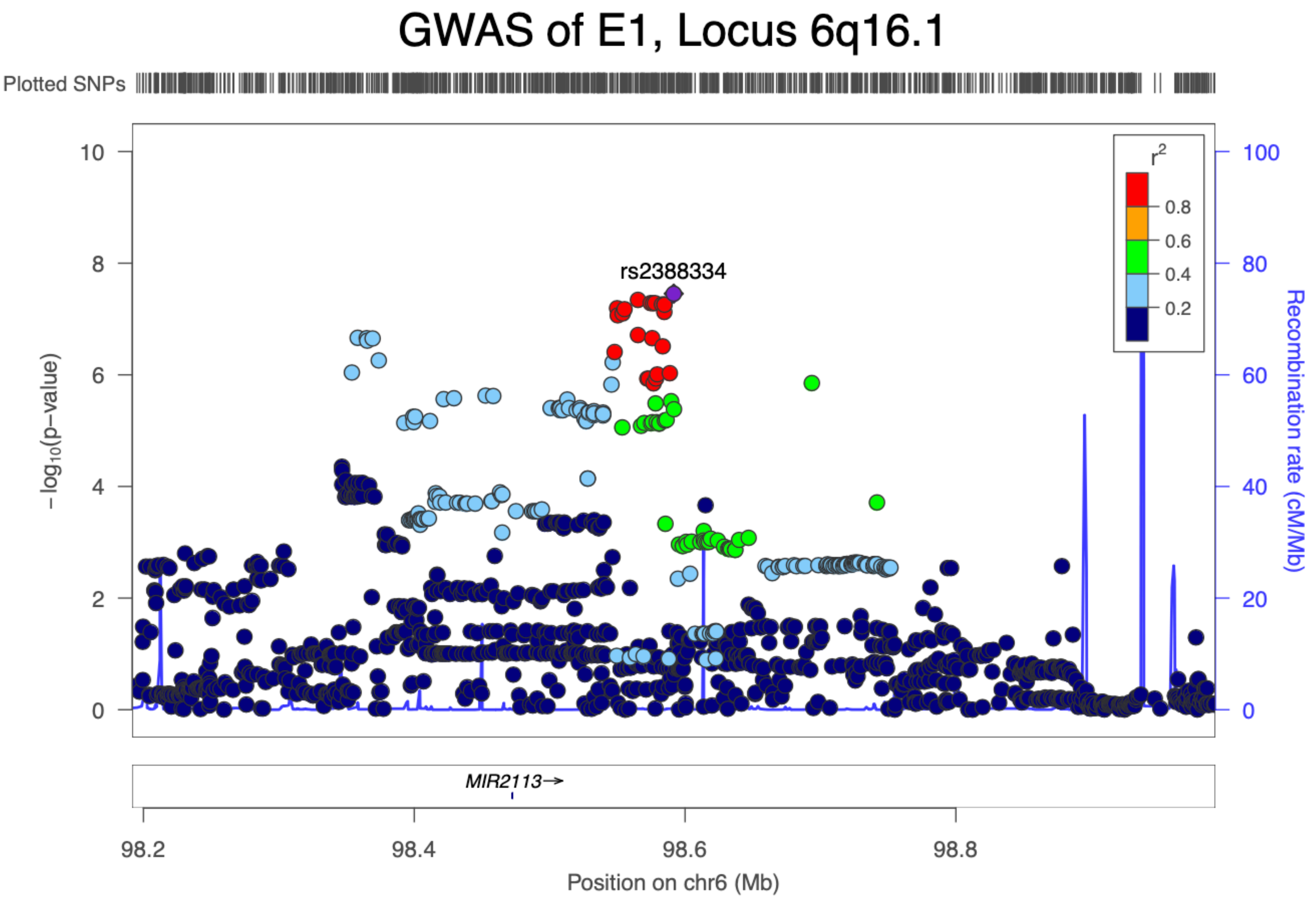
GWAS of E1, Locus 6q16.1
Plotted SNPs
rs2388334
r²
0.8
0.6
0.4
0.2
−log10(p−value)
Recombination rate (cM/Mb)
MIR2113→
98.2
98.4
98.6
98.8
Position on chr6 (Mb)
0
2
4
6
8
10
20
40
60
80
100

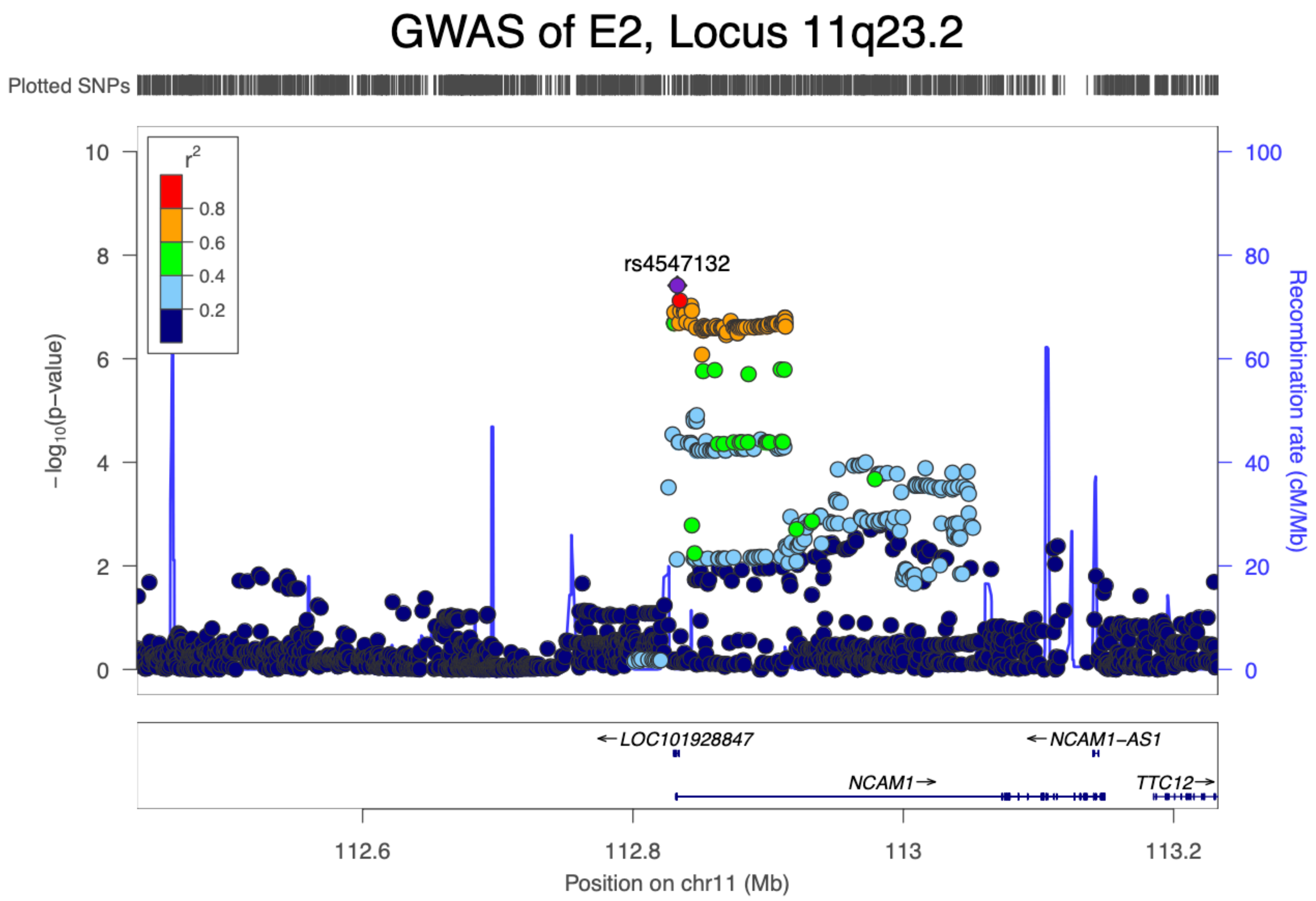
GWAS of E2, Locus 11q23.2
Plotted SNPs
r2
0.8
0.6
0.4
0.2
rs4547132
−log10(p−value)
Recombination rate (cM/Mb)
10
8
6
4
2
0
100
80
60
40
20
0
←LOC101928847
←NCAM1−AS1
NCAM1→
TTC12→
112.6
112.8
113
113.2
Position on chr11 (Mb)

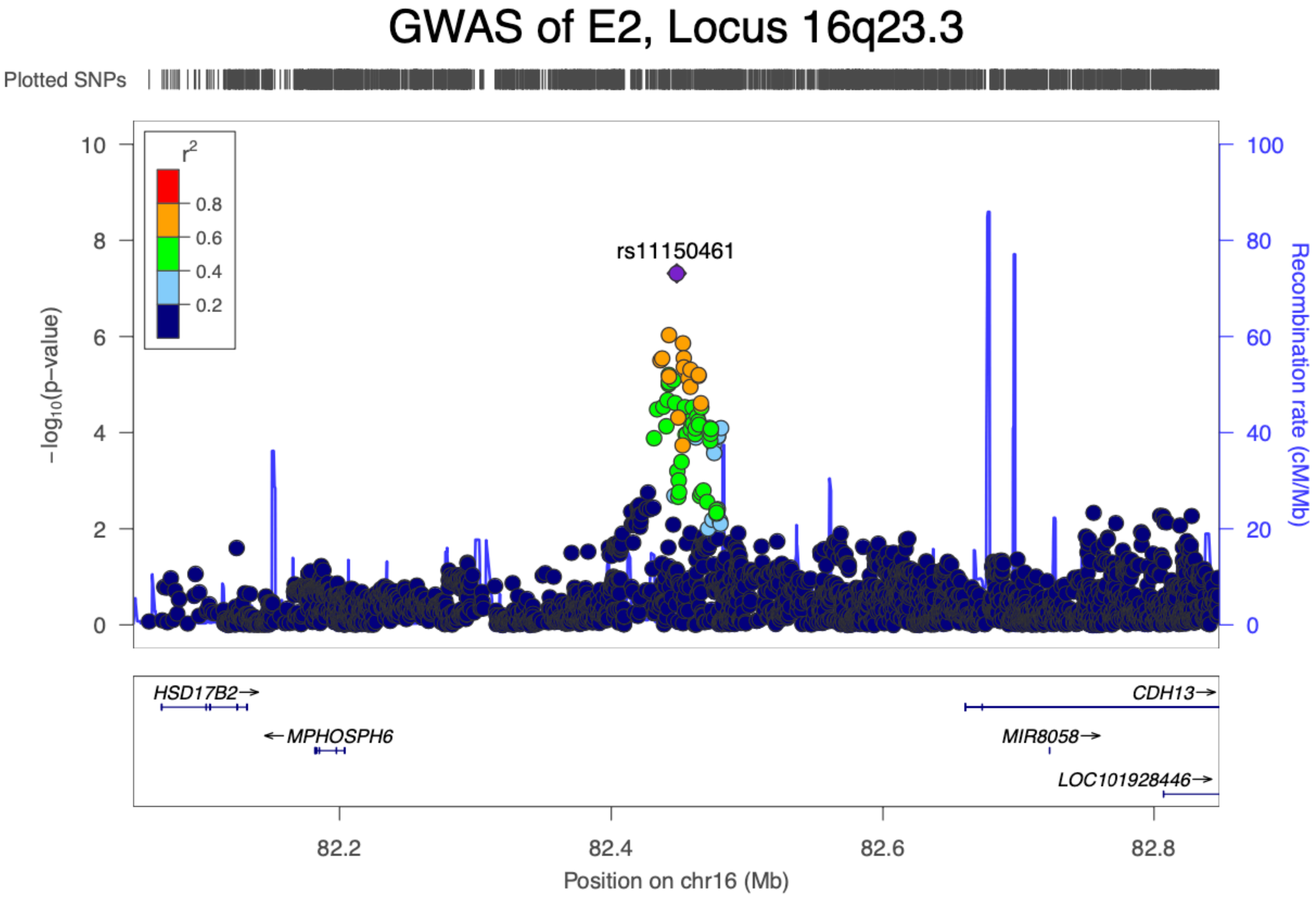

GWAS of E2, Locus 16q23.3
Plotted SNPs
r2
0.8
0.6
0.4
0.2
rs11150461
-log10(p-value)
Recombination rate (cM/Mb)
HSD17B2
MPHOSPH6
CDH13
MIR8058
LOC101928446
82.2
82.4
82.6
82.8
Position on chr16 (Mb)

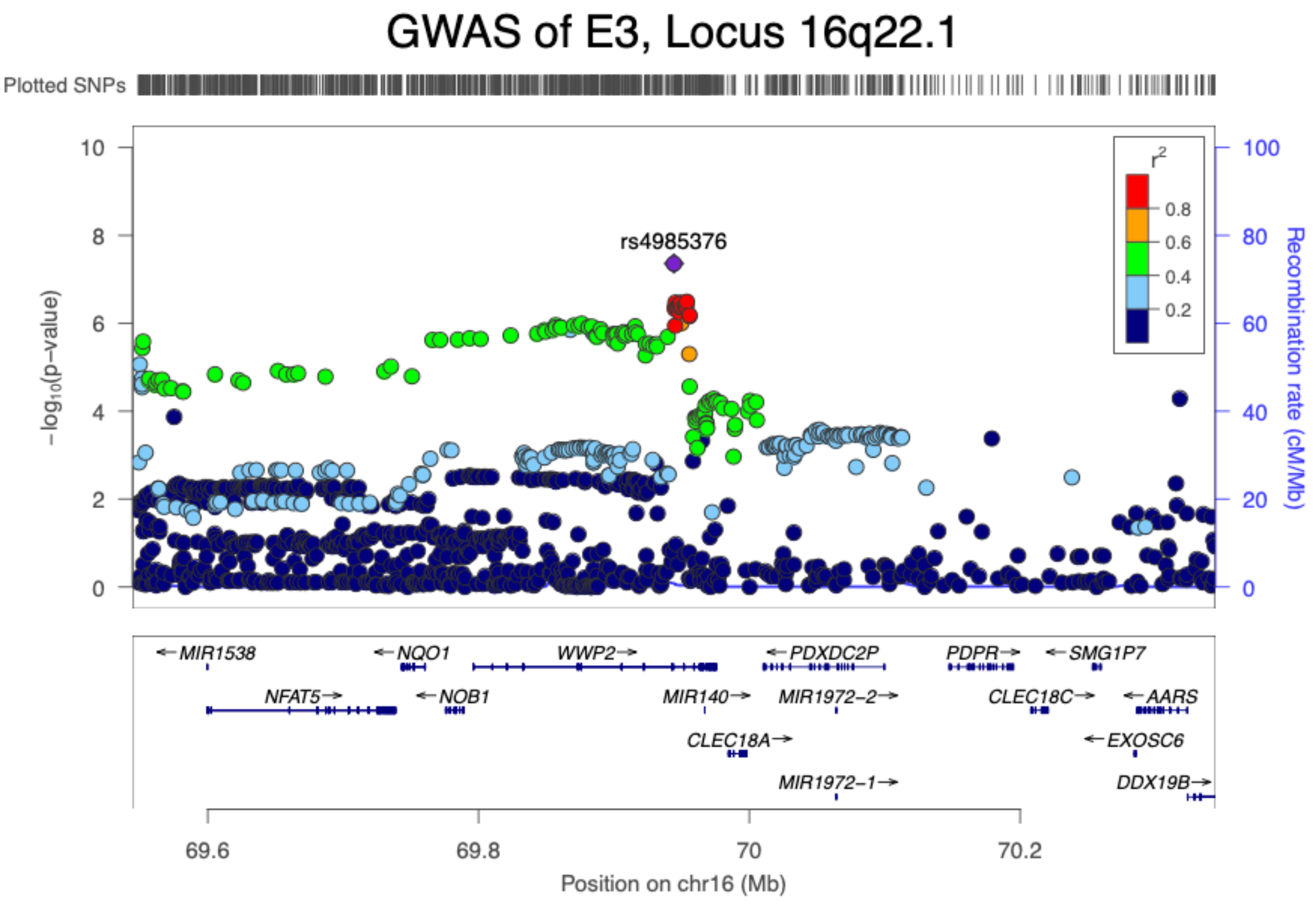
GWAS of E3, Locus 16q22.1
Plotted SNPs
rs4985376
r2
0.8
0.6
0.4
0.2
-log10(p-value)
Recombination rate (cM/Mb)
10
8
6
4
2
0
100
80
60
40
20
0
MIR1538
NQO1
WWP2
PDXDC2P
PDPR
SMG1P7
NFAT5
NOB1
MIR140
MIR1972-2
CLEC18C
AARS
CLEC18A
EXOSC6
MIR1972-1
DDX19B
69.6
69.8
70
70.2
Position on chr16 (Mb)
