## Supplementary Dataset 2 for "Genome-wide association study of school grades identifies a genetic overlap between language ability, psychopathology and creativity"

**Supplementary dataset 2 | Phenome-wide association plots.**

The PheWAS plots of seven variants are in this file. The seven variants are index variants in the genome-wide loci identified in this study--four for E1, two for E2 and one for E3. The PheWAS was done using GWASatlas online tool (atlas.ctglab.nl). Totally 4756 phenotypes were included. Only associations that are statistically significant after multiple testing correction ($P<1.05\times10^{-5}$) were shown in the plots. Not all points can be labelled. Hence, only relevant phenotypes or phenotypes considered as interesting were labelled.

**Phenotype:** E1
**Variant ID:** rs7613360

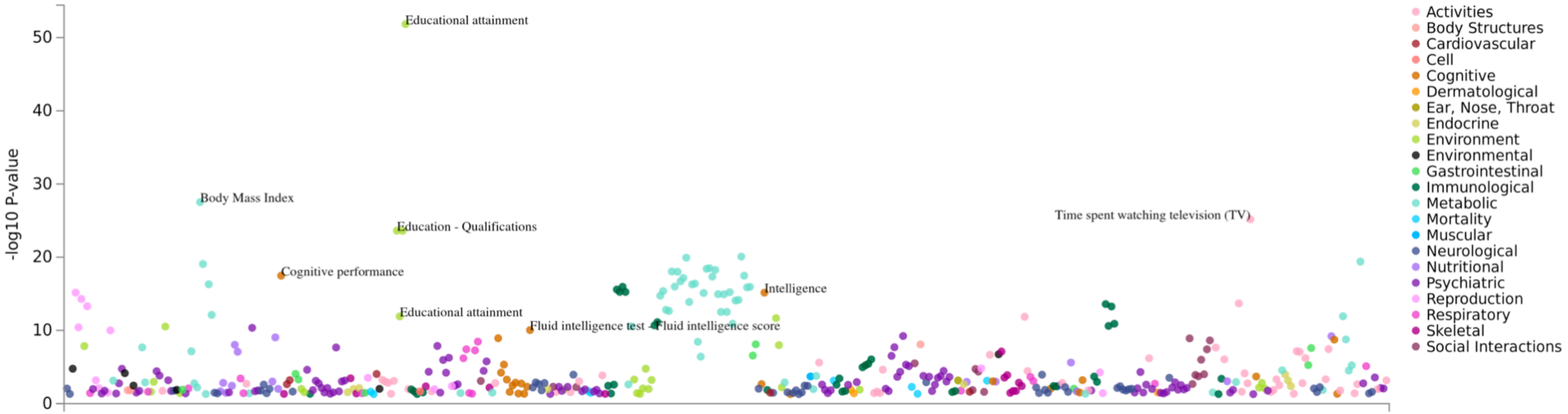

**Phenotype:** E1
**Variant ID:** rs2388334

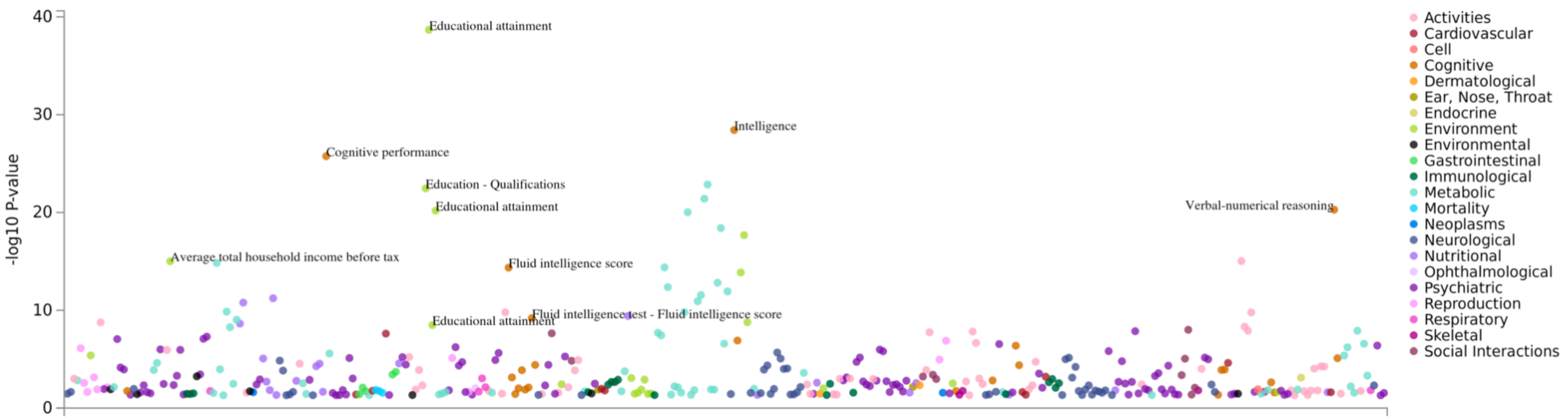

**Phenotype:** E1
**Variant ID:** rs11895772

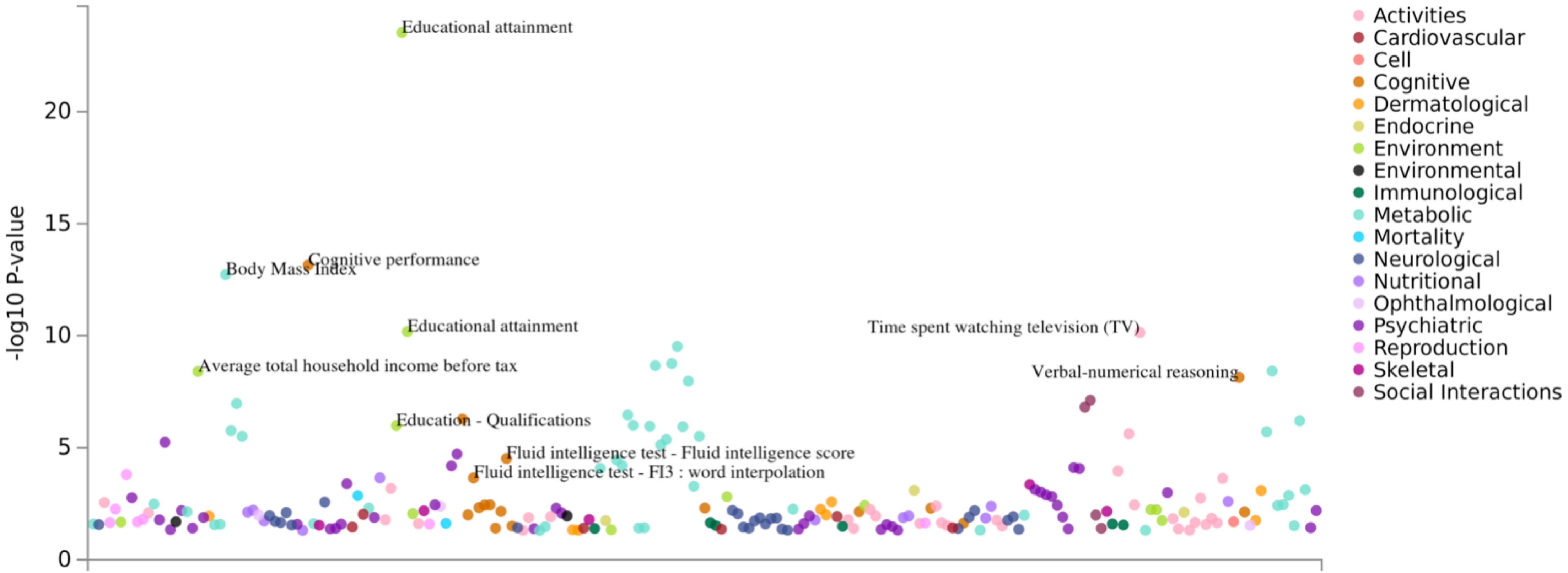

**Phenotype:** E1
**Variant ID:** rs696732

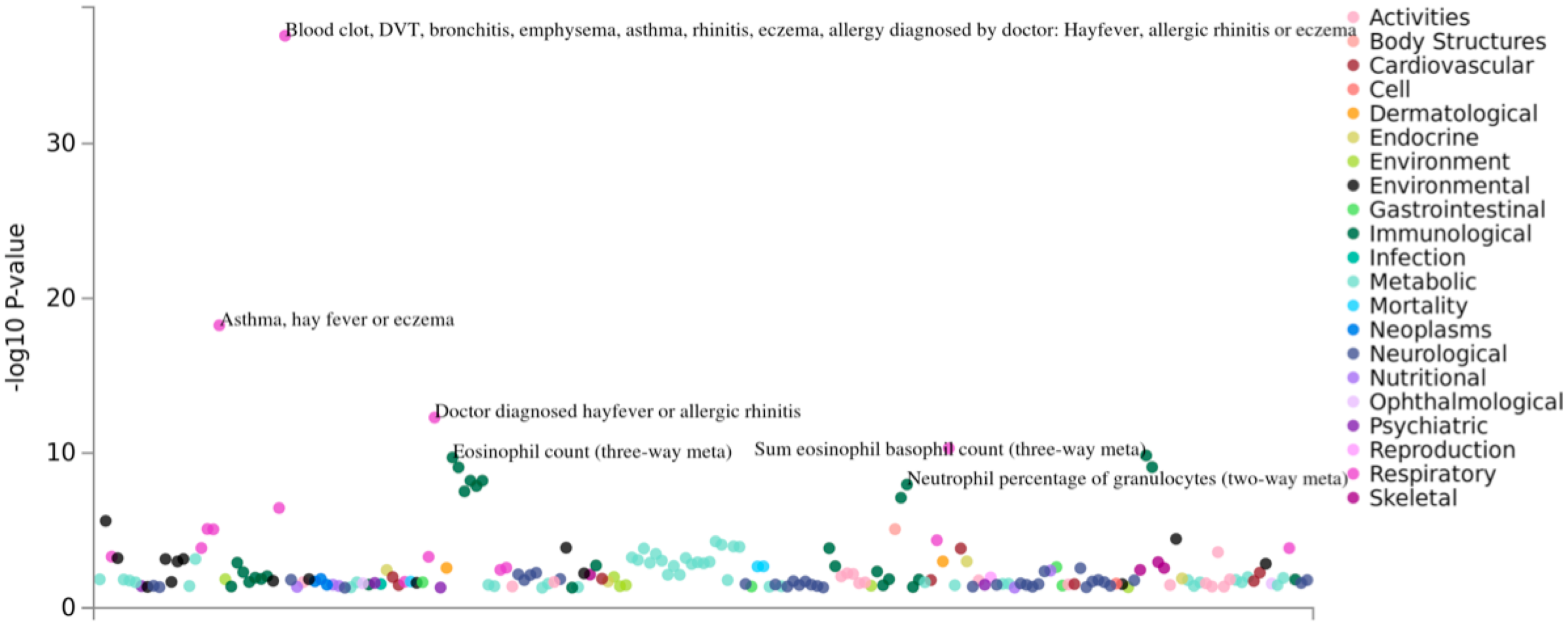

**Phenotype:** E2
**Variant ID:** rs4547132

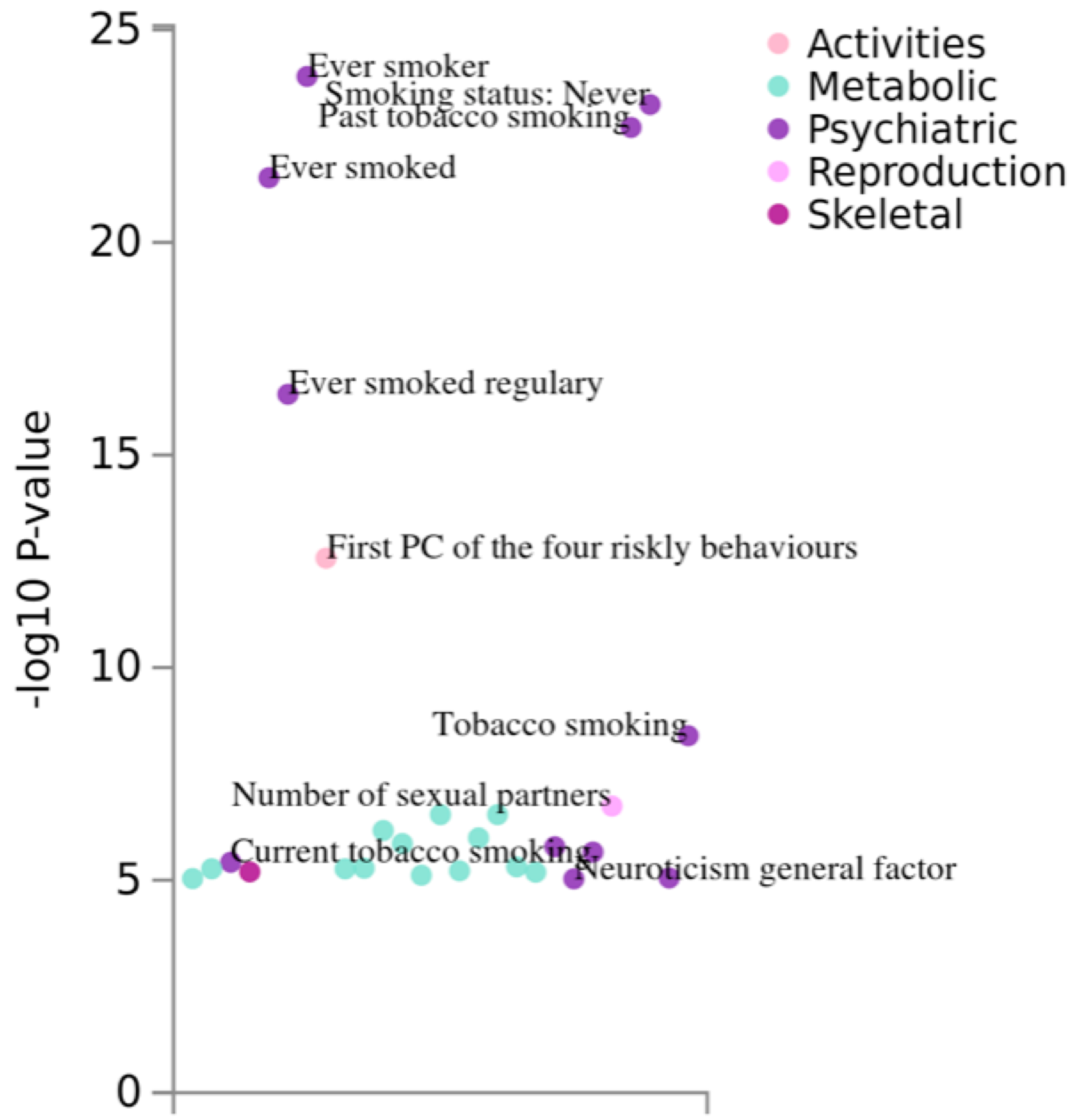

**Phenotype:** E2
**Variant ID:** rs11150461

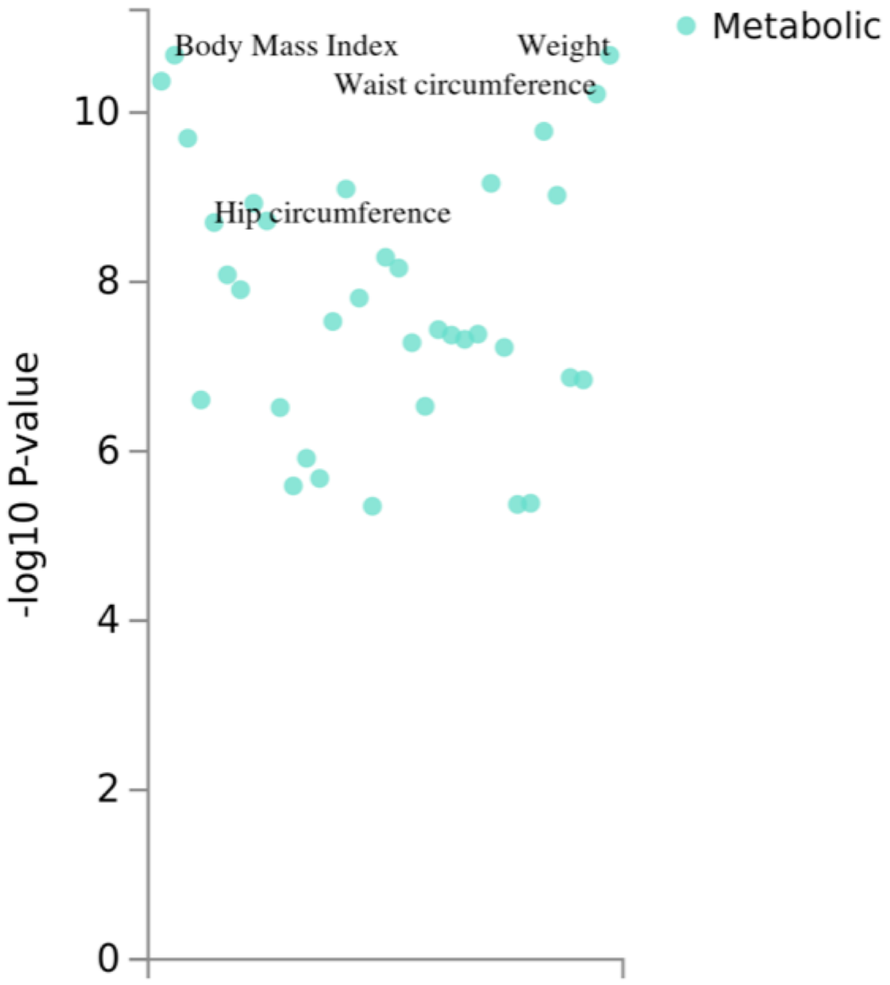

**Phenotype:** E3
**Variant ID:** rs696732

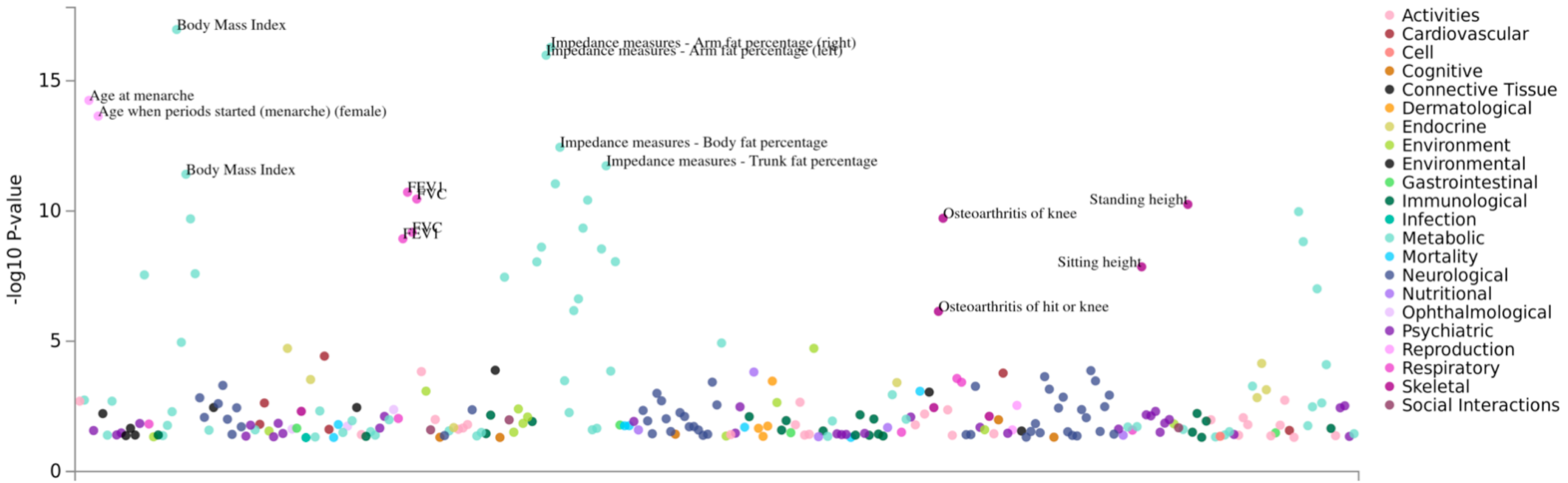
